# Dopamine shifts striatal ramping dynamics

**DOI:** 10.64898/2026.11.25.769161

**Authors:** Alexandra S. Bova, Rachael A. Volkman, Vaibhav Khandelwal, Matthew A. Weber, Madison McMurrin, Julia Kryca, Kartik Sivakumar, Christopher M. Hunter, Christina M. Weber, Jeffrey D. Dawson, Ergun Y. Uc, Nandakumar S. Narayanan

## Abstract

Diseases that disrupt dopamine impair cognition; however, it is unknown how dopamine affects neuronal networks that instantiate cognitive processing. We studied this problem in the dorsomedial striatum during interval timing, an elementary cognitive task requiring working memory and attention to estimate an interval of several seconds. Interval timing requires medium spiny neurons (MSNs) that ramp up or down over temporal intervals. While these neurons receive prominent dopaminergic input from the substantia nigra, it is unknown how dopamine modulates MSN activity during interval timing. Here, we report four new findings. First, ramping activity of MSN ensembles predicts interval timing behavior. Second, dopamine dynamics predict timing behavior and preferentially predict the slope of MSN ensembles that ramp down; these relationships are not observed during tasks without temporal control such as Pavlovian conditioning. Third, inactivating substantia nigra dopamine disrupts timing as well as MSNs that ramp down, but not those that ramp up. Fourth, a drift-diffusion model perturbing down-ramping accurately predicted interval timing behavior in mice with dopamine inhibition and in humans with Parkinson’s disease. Together, these results reveal how dopamine shapes striatal ramping dynamics and suggest that disrupting dopamine can alter specific trajectories of cognitive computations. These data are relevant for understanding the cognitive striatum and for generating new markers and therapeutic strategies for human striatal and dopaminergic diseases.

## INTRODUCTION

Dopamine is a multifaceted neurotransmitter with roles in movement, motivation, reward prediction error, learning, and salience^1–5^. One common and debilitating feature of diseases affecting dopamine is cognitive dysfunction. For instance, in Parkinson’s disease, cognitive impairments are major drivers of morbidity and mortality, but there are few reliable treatments^6^. Other diseases disrupting dopamine, such as attention deficit-hyperactivity disorder and schizophrenia, also feature prominent cognitive dysfunction that lacks effective treatments^7,8^. Thus, there is an urgent need to determine mechanisms of how dopamine impacts cognitive processing to develop better diagnostic and treatment strategies.

Cognitive function is complex and challenging to study in mechanistic mouse models in which dopamine and neuronal activity can be measured in detail. One elementary cognitive paradigm that provides traction on this problem is interval timing, which requires participants to estimate intervals of several seconds by making a motor response. Interval timing relies on cognitive functions such as working memory for temporal rules and attention to the passage of time^9^. Importantly, interval timing is a cognitive paradigm that translates across species from mouse models to human disease^10–13^. Both dopamine and striatal medium spiny neurons (MSNs) play critical roles during interval timing tasks, with dopamine disruptions reliably slowing timing behavior^11,14,15,15,16^ and striatal MSNs representing temporal information across intervals^3,15,17–22^. MSNs encode time via ramping activity, or monotonic changes in firing rate over the interval, with D1 MSNs tending to ramp down, and D2 MSNs tending to ramp up^15^. Despite this body of work, it is unknown how dopamine affects MSN ramping.

We investigated this problem by recording striatal MSNs while simultaneously recording from and manipulating dopamine during interval timing to test the hypothesis that dopamine affects striatal ramping activity. We find evidence that (1) MSN ensemble ramping activity predicts interval timing; (2) dopamine dynamics preferentially track the slope of down-ramping MSN ensembles during timing but not in tasks without temporal control, such as Pavlovian conditioning; (3) dopaminergic inactivation selectively disrupts timing behavior and down-ramping MSNs; and (4) drift-diffusion models of down-ramping successfully predict timing deficits in both mice and humans with Parkinson’s disease. Together, these insights reveal how dopamine modulates striatal dynamics during cognitive computations, providing new perspective on the cognitive striatum and associated neurodegenerative disorders.

## RESULTS

### Dorsomedial striatal medium spiny neuron ensembles predict time

To interrogate striatal dopaminergic circuits during cognitive processes, we turned to interval timing, which requires participants to estimate an interval of several seconds by making a motor response. We leveraged a mouse-optimized interval timing task in which mice switch responding between nosepokes to indicate that an interval of 6 seconds has elapsed, as described in depth in prior work (Fig. 1A)^15,22–25^. Briefly, trials begin with a response at the back nosepoke, which initiates both auditory and visual cues. On 50% of trials, cues sustain for 6 seconds and the first response at the short nosepoke after 6 seconds is rewarded; these trials are not analyzed. On the other 50% of trials, cues sustain for 18 seconds and responses at the long nosepoke after 18 seconds trigger rewards. Therefore, mice must switch responding from the short to the long nosepoke after estimating that 6 seconds has elapsed. The time that mice depart the short nosepoke before arriving at the long nosepoke is defined as the *switch response time*. Because cues are identical on all trials, the switch response time is a time-based decision that reflects the mouse’s internal estimate of elapsed time. As in prior studies^15,22,24^, switch responses peaked after 6 seconds, just after short nosepoke responses peaked, and long nosepokes peaked as mice approached trial end (Fig. 1B and Extended Data Fig. 1A). Similarly, movement tracking with DeepLabCut^26^ revealed that mice consistently moved quickly from the back nosepoke to the short nosepoke, then switched to responding at the long nosepoke (Extended Data Fig. 1B-C).

**Figure 1.**
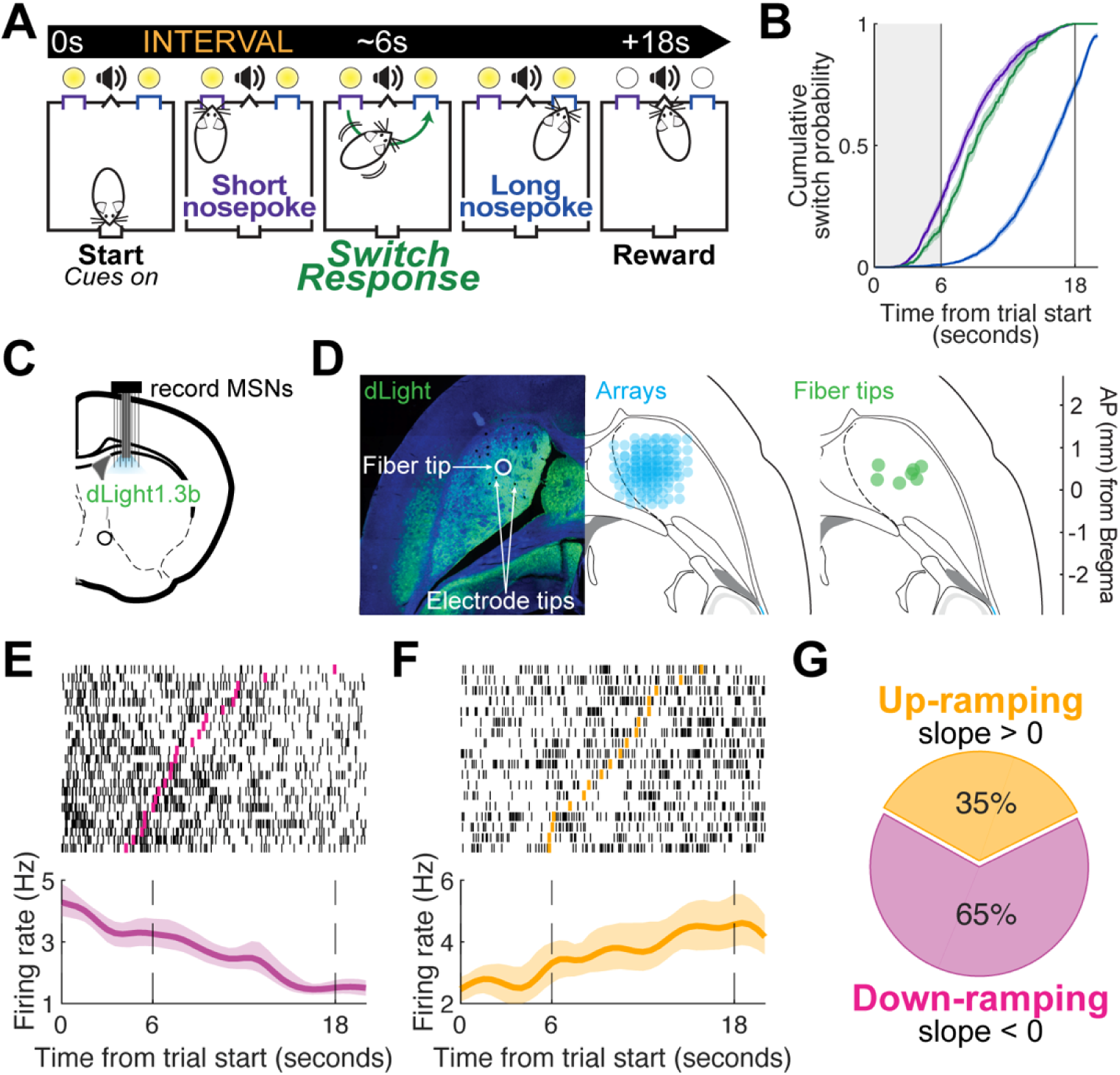
Mouse-optimized interval timing. **A)** We trained mice in an interval timing task in which they indicate their estimation of time by switching between nosepokes. Trials begin when mice respond at a back nosepoke, which initiates visual and auditory cues. On 50% of trials, cues sustain for 6 seconds and mice earn a reward for the first response at the short nosepoke after 6 seconds. On the other 50% of trials, cues sustain for 18 seconds and mice begin responding at the short nosepoke, but *switch* responding to the long nosepoke when they estimate that 6 seconds have elapsed. Mice are rewarded for the first response at the long nosepoke after 18 seconds. **B)** Cumulative response density from 494 trials in 25 mice (13 female) for short nosepoke (purple), long nosepoke (blue), and switch responses (green). Shading represents s.e.m. **C)** We recorded dLight1.3b fiber photometry simultaneously with electrophysiological recordings of medium spiny neuron (MSN) ensembles in the DMS. **D)** Left: Horizontal section with dLight expression (green). Circle identifies fiber optic tip and arrows identify electrode tips. Right: Horizontal sections depicting electrode reconstruction (blue) and fiber optic locations (green). **E)** Peri-event raster (top) and estimated firing rate (bottom) of an individual MSN that ramps down and **F)** of an individual MSN that ramps up across the timed interval. Pink and yellow ticks represent switch response times. Shading represents bootstrapped 95% confidence interval. **G)** Percentage of neurons that have positive slopes (“up-ramping”, yellow) and negative slopes (“down-ramping”, pink). Values in pie charts are rounded to the nearest whole number. Data from 104 MSNs and 314 trials in 8 mice.

In 8 mice (Supplemental Table 1), we recorded ensembles of medium spiny neurons (MSNs) with multi-channel electrode arrays (Fig. 1C-D). As previously reported^15,22,25,27,28^, MSNs displayed ramping activity, defined as a monotonic change in firing rate across the 18-second interval (Fig 1E-F). We classified individual MSNs according to the direction of their GLM-derived slope estimates from 0 to 18 seconds. Neurons with negative slopes were classified as ‘down-ramping’ (Fig. 1E, G; 65.4% of neurons; Median slope: -0.26 (IQR: -0.49 to -0.11)), whereas neurons with positive slopes were classified as ‘up-ramping’ (Fig. 1F, G; 34.6% of neurons; Median slope: 0.52 (IQR: 0.13 to 0.73)).

Past work has extensively described how MSN ramping ensembles can encode elapsed time^15,17,18,20,22^. We compared how down-ramping MSN vs. up-ramping MSN ensembles predicted elapsed time using naïve Bayesian classifiers as in our past work^10,20,22^. This approach decoded trial-by-trial predictions of time from neuronal ensemble firing rates and quantified predictions via R^2^ of objective time vs. predictive time for each trial (Fig 2A-B). Temporal decoding was similar between ensembles of all MSNs, down-ramping MSNs, and up-ramping MSNs for both the full trial (0 – 18 seconds; all vs. down-ramping: *p* = 0.99; all vs. up-ramping: *p* = 0.39; down-ramping vs. up-ramping: *p* = 0.67) and from 0 to 6 seconds (all vs. down-ramping: *p* = 1.0; all vs. up-ramping: *p* = 0.67; down-ramping vs. up-ramping: *p* = 0.74). These results are in line with past work demonstrating that MSN ensembles encode elapsed time.

**Figure 2.**
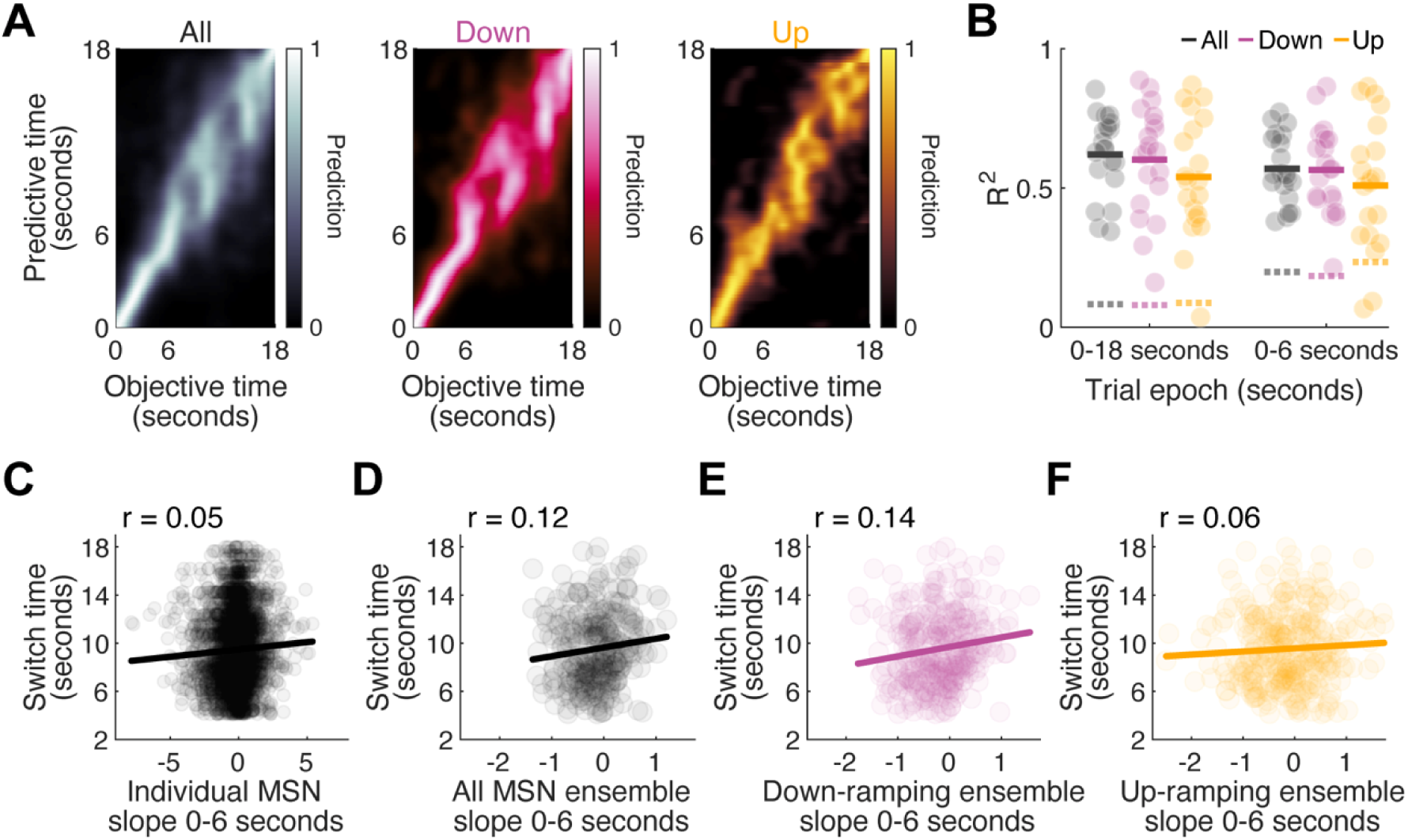
Striatal MSN ensembles predict time. **A)** We trained a naïve Bayesian classifier to decode temporal predictions on a trial-by-trial basis from firing rate from the entire MSN ensemble (gray), an ensemble of only down-ramping MSNs (pink), and an ensemble of only up-ramping MSNs (yellow). **B)** A naïve Bayesian classifier found that all ensemble classes (all MSNs, down-ramping MSNs, up-ramping MSNs) decoded time similarly for 0 to 18 seconds and for 0 to 6 seconds. Each dot represents a trial and solid horizontal lines represent trial averages. Dashed horizontal lines represent trial averages for shuffled data. **C)** The slope from 0 to 6 seconds of individual MSNs weakly correlated with switch response time. 3904 data points from 104 MSNs in 8 mice. **D)** The slope from 0 to 6 seconds of all MSN ensembles predicted switch response time. Each dot represents an individual trial. Data from 300 trials in 104 MSNs in 8 mice. **E)** The slope from 0 to 6 seconds of down-ramping MSN ensembles predicted switch response time. Each dot represents an individual trial. Data from 300 trials in 68 MSNs in 8 mice. **F)** The slope from 0 to 6 seconds of up-ramping MSN ensembles did not predict switch response time. Each dot represents an individual trial. Data from 300 trials in 36 MSNs in 8 mice.

If ramping activity was important for temporal control of action, the slope should predict animals’ internal estimates of time. We examined regression-based slopes of MSN activity from 0 to 6 seconds, and animals’ switch response time, which reflects the moment when they estimate at least 6 seconds have elapsed and they switch from the short to the long nosepoke. The slope of individual MSNs was a significant – but weak – predictor of switch response time (Fig 2C, r = 0.05, *p* = 0.003) and the average slope across an individual mouse’s ensemble of MSNs also predicted switch response time (Fig 2D, r = 0.12, *p* = 0.05). We discovered that down-ramping MSN ensembles predicted switch response time (Fig 2E, r = 0.14, *p* = 0.01) while up-ramping MSN ensembles did not (Fig 2F, r = 0.06, *p* = 0.30), implying specific ramping dynamics were linked with temporal control of action. Together, these data demonstrate that while all MSNs encode elapsed time, down-ramping MSN ensembles preferentially predicted temporal control of action.

### Dorsomedial striatal dopamine predicts timing behavior and down-ramping MSN ensembles

To quantify dopamine dynamics alongside dorsomedial striatal MSN activity, we recorded fiber photometry with the fluorescent dopamine sensor dLight1.3b^31^. As expected, DMS dLight activity was strongly decreased with SNc dopamine neuron inhibition for both 2 and 7 seconds (Fig. 3A-B; *n* = 2; AUC laser off vs. laser on 2 seconds: *p* < 0.0001; 7 seconds: *p* < 0.0001). To investigate DMS dopamine dynamics, we recorded dLight in a separate cohort of 8 well-trained mice during interval timing (Fig. 3C). We found prominent modulations in dopamine dynamics, as measured by dLight z-score, with dLight z-score elevated at trial start and reward delivery (Fig. 3D-E and Extended Data Fig. 2A-B). dLight signal decreased prominently before the switch response but the amplitude of this signal did not predict response time (-4 to 0 seconds prior to switch response; Fig 3G-H, r = 0.02, *p* = 0.77). However, the slope of the dLight signal reliably predicted switch response time with a more negative slope predicting an earlier switch response (Fig 3I; r = 0.24, *p* < 0.0001). Importantly, the dLight signal around non-switch response short nosepoke exits was distinct from the dLight signal around switch responses and did not predict switch response time (Extended Data Fig. 2C-F).

**Figure 3.**
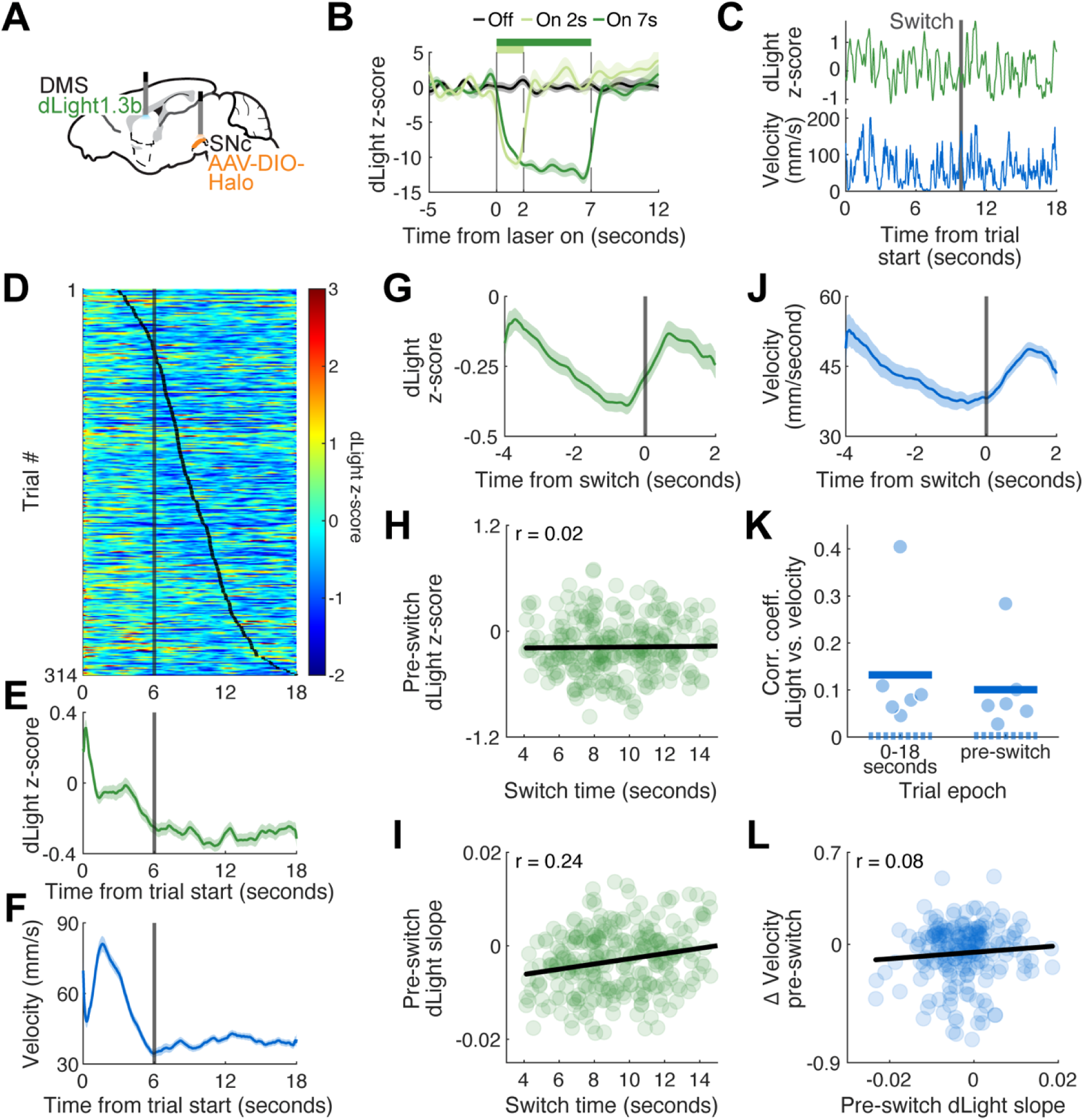
Dorsomedial striatal dopamine predicts interval timing. **A)** We injected Cre-dependent Halorhodopsin into the substantia nigra pars compacta (SNc) and dLight1.3b into the dorsomedial striatal (DMS) of 2 DAT^IRES^-Cre mice. **B)** Laser light (594 nm, 2 mW) randomly delivered for 2 or 7 seconds into fiber optics implanted above the SNc decreased dLight signal in the DMS. Data from 21 Off trials, seven 2s On trials, and twelve 7s On trials in 2 mice. **C)** Top: dLight z-score from a single switch response trial. Bottom: Movement velocity (mm/second) during the same switch response trial. Solid gray line represents the switch response time. **D)** dLight z-score during every switch response trial. Trials are sorted by switch response time (indicated by black tick marks) and aligned to trial start. Data from 314 trials in 8 mice. **E)** Average dLight z-score across all switch response trials. Data from 314 trials in 8 mice. **F)** Average movement velocity (mm/second) from all switch response trials. Data from 212 trials in 6 mice with motion tracking. **G)** Average dLight z-score aligned to the switch response. Data from 314 trials in 8 mice. **H)** The average dLight z-score in the four second period prior to the switch response did not predict switch response time. Each dot represents a single trial. Data from 273 trials in 8 mice; 27 outliers removed. **I)** The slope of the dLight signal in the four second period prior to the switch response predicted switch response time. Each dot represents a single trial. Data from 276 trials in 8 mice; 24 outliers removed. **J)** Average movement velocity (mm/second) aligned to the switch response. Data from 212 trials in 6 mice. **K)** Pearson’s correlation coefficients for each individual mouse between moment-by-moment dLight signal (z-score) and movement velocity (mm/second) across the full trial interval (0 – 18 seconds) and in the four seconds preceding the switch response (‘pre-switch’). Horizontal lines represent group averages. Dashed lines represent group averages for shuffled data. Data from 212 trials in 6 mice. **L)** Δ velocity in the four second period preceding the switch response did not predict the slope of the dLight signal in this period. Each dot represents a single trial. Data from 191 trials in 6 mice; 8 outliers removed. Shading in B, E, F, G, and J represents s.e.m.

One important confound is movement, as mice move throughout the task from the back to the short nosepoke, and when switching from the short to the long nosepoke (Fig 3F, J). Velocity was somewhat predictive of dLight z-score across the full trial (average r = 0.13; *p* = 0.002 vs. shuffled data) and in the pre-switch period (average r = 0.10; *p* = 0.002 vs. shuffled data; Fig 3K). However, pre-switch Δvelocity and pre-switch dLight slope were not correlated (Fig 3L, r = 0.08, *p* = 0.26). To further control for movement velocity’s contribution to the dLight slope-switch time relationship, we ran a partial correlation of dLight slope and switch time accounting for velocity. Indeed, dLight slope and switch time were strongly correlated (r = 0.19, *p* = 8.95×10^-3^ in 5 mice with dLight recordings and velocity tracking) and this relationship remained significant when partial correlation controlled for velocity (r = 0.16, *p* = 0.03). Including velocity did not substantially affect the relationship between dopamine and switch response time (comparison of correlation coefficient of correlation vs. partial correlation via Steiger Z-test: *p* = 0.44). Therefore, although movement velocity contributes to dopamine dynamics, it cannot fully explain the predictive relationship of dLight slope on switch response time.

Next, we interrogated the relationship between dopamine dynamics and MSN ensembles. We recorded dopamine dynamics with dLight fiber photometry simultaneously with MSN ensemble recordings and motion tracking (Fig. 4A). We first compared decoding of MSN ensembles and dLight dynamics using naïve Bayesian classifiers (Fig. 4B). Temporal decoding was stronger in MSN ensembles compared to temporal decoding in dLight dynamics across the full trial (Fig. 4C, 0 – 18 seconds; average R^2^ MSNs = 0.28, average R^2^ dLight = 0.06; *p* < 0.0001) and from 0 to 6 seconds (average R^2^ MSNs = 0.38, average R^2^ dLight = 0.06; *p* < 0.0001). These data provide remarkable evidence that while MSN ensembles encode elapsed time, dopamine dynamics do not.

**Figure 4.**
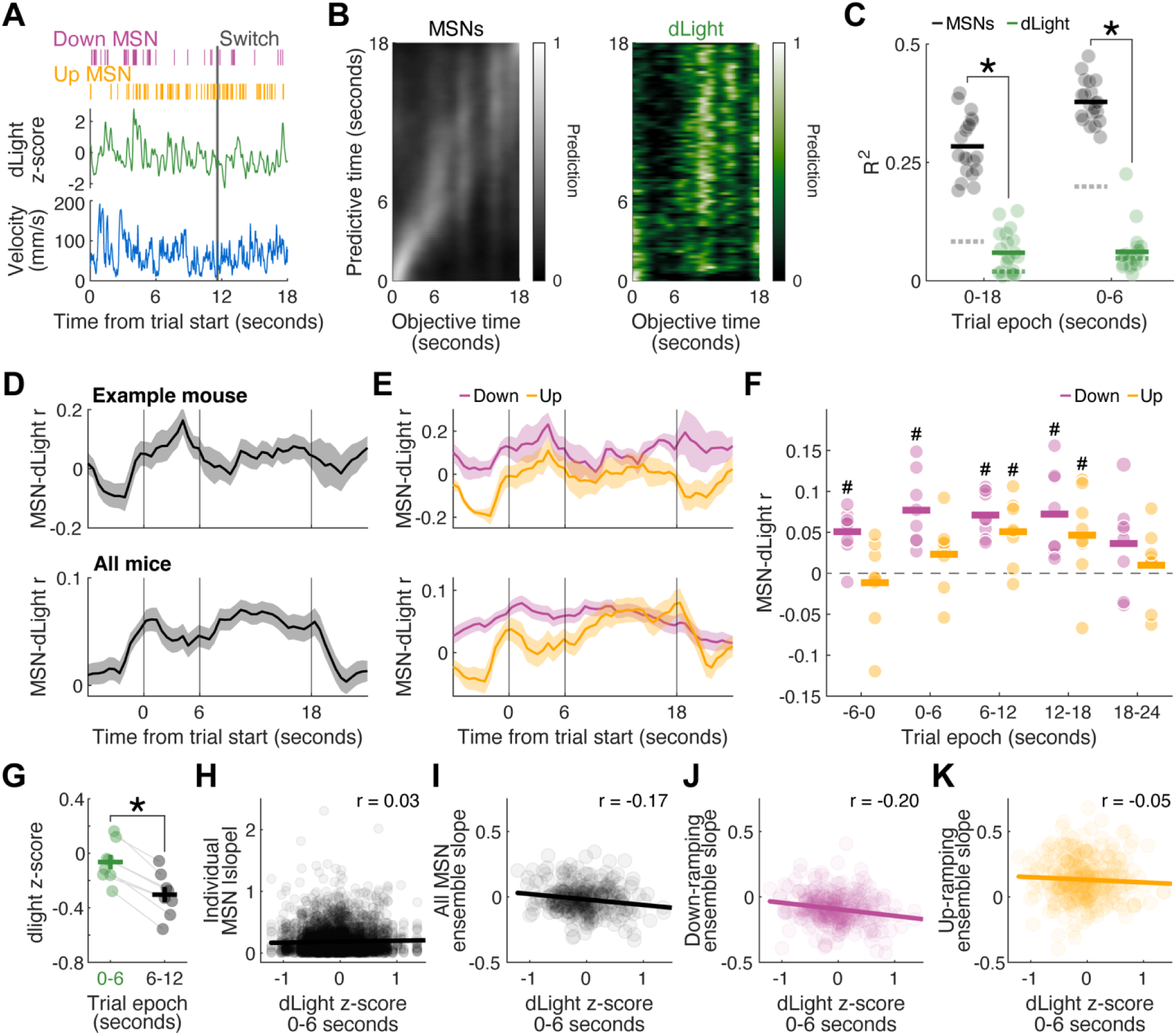
Striatal dopamine predicts down-ramping MSN ensemble slope. **A)** Top: Peri-event raster from a single switch response trial from a down-ramping MSN (pink) and an up-ramping MSN (yellow). Middle: dLight signal (z-score) during the same switch response trial. Bottom: Movement velocity (mm/second) during the same switch response trial. Solid gray line represents the time of the switch response. **B)** We trained naïve Bayesian classifiers to make temporal predictions on a trial-by-trial basis from firing rate of MSN ensembles (gray) and from dLight signal (green). **C)** Naïve Bayesian classifiers found that MSN ensembles decoded time more accurately than dLight signal across both 0 to 18 seconds and 0 to 6 seconds. Each dot represents a temporal decoding trial and solid horizontal lines represent averages. Dashed horizontal lines represent averages for shuffled data. Data from 314 trials in 8 mice. **D)** Average moment-by-moment correlation between dLight and individual MSNs during switch trials for a single mouse (top) and averaged across all mice (bottom). Data from 314 trials in 8 mice. **E)** Average correlation between dLight and individual down-ramping MSNs (pink) or between dLight and individual up-ramping MSNs (yellow) during switch trials for a single mouse (top) and averaged across all mice (bottom). Data from 314 trials in 8 mice. **F)** dLight signal was correlated with down-ramping MSNs from -6 to 0 seconds, 0 to 6 seconds, 6 to 12 seconds, and 12 to 18 seconds. dLight signal was correlated with up-ramping MSNs only from 6 to 12 and 12 to 18 seconds. Data from 314 trials in 8 mice. **G)** The average dLight signal was significantly higher from 0 to 6 seconds than from 6 to 12 seconds. Horizontal lines in F-G represent group average, vertical lines represent s.e.m., and dots represent individual mouse average. Data from 271 trials in 8 mice. **H)** The average dLight signal from 0 to 6 seconds did not predict |slope| of individual MSNs. 4106 data points from 104 MSNs in 8 mice. **I)** The average dLight signal from 0 to 6 seconds predicted the average ensemble slope from all MSNs. Data from 314 trials in 104 MSNs in 8 mice. **J)** The average dLight signal from 0 to 6 seconds predicted the average ensemble slope from down-ramping MSNs. Data from 314 trials in 68 MSNs in 8 mice. **K)** The average dLight signal from 0 to 6 seconds did not predict the average ensemble slope from up-ramping MSNs. Data from 314 trials in 36 MSNs in 8 mice. * *p* < 0.05, *# p* < 0.05 compared to zero

Corticostriatal input regulates striatal MSN ramping activity^32^, but it is unknown how striatal dopamine release contributes to MSN ramping. We first investigated if dLight dynamics predicted MSN firing rate activity during switch trials (Fig. 4D). dLight z-score was correlated with down-ramping MSNs before trial start and during the trial, but not after trial end (Fig. 4E-F, average r vs. 0: -6-0 seconds: *p* = 0.01; 0-6 seconds: *p* < 0.0001; 6-12 seconds: *p* = 1.3×10^-4^; 12-18 seconds: *p* = 1.0×10^-4^; 18-24 seconds: *p* = 0.08). Interestingly, up-ramping MSNs were only correlated with dLight signal late in the trial (average r vs. 0: -6-0 seconds: *p* = 0.92; 0-6 seconds: *p* = 0.40; 6-12 seconds: *p* = 0.01; 12-18 seconds: *p* = 0.02; 18-24 seconds: *p* = 0.92).

Next, we asked if dLight dynamics early in the trial (0-6 seconds or pre-switch response; dLight 0-6 seconds vs. dLight 6-12 seconds: *p* < 0.0001) correlated with MSN ramping dynamics (Fig. 4G and Extended Data Fig. 3A). Early trial dLight z-score weakly predicted individual MSN slope (Fig. 4H and Extended Data Fig. 3B; r = 0.03, *p* = 0.03). Strikingly, dLight z-score negatively predicted the average MSN ensemble slope such that higher dopamine was associated with a more negative slope (Fig. 4I and Extended Data Fig. 3C, r = -0.17, *p* = 0.002). This pattern was driven by down-ramping MSN ensembles (Fig. 4J and Extended Data Fig. 3D, r = -0.20, *p* = 4.72×10^-4^), but not by up-ramping MSN ensembles (Fig. 4K and Extended Data Fig. 3E, r = -0.05, *p* = 0.33). Collectively, these results provide novel evidence that striatal dopamine dynamics early in the trial regulate down-ramping MSN ensembles but not up-ramping MSN ensembles.

### Time-related dopamine and MSN modulations do not reflect anticipation of reward

Both striatal dopamine and MSN activity predicted temporal control of action. However, it is possible that this modulation reflects anticipation of upcoming reward unrelated to time. To disambiguate reward anticipation versus interval timing, we recorded dLight and MSN activity from the same mice (*n* = 5 out of 8 mice from Fig. 4) during Pavlovian conditioning (Fig. 5A and Extended Data Fig. 4A). Importantly, during Pavlovian conditioning cues predict reward delivery after a fixed time interval. Therefore, mice can learn to anticipate reward delivery based on the passage of time but are not required to control movements in time or actively track the passage of time to earn a reward. Due to shorter session times, average reward rate was higher during Pavlovian conditioning sessions, but mice earned similar numbers of rewards across the two sessions (Extended Data Fig. 4B-C).

**Figure 5.**
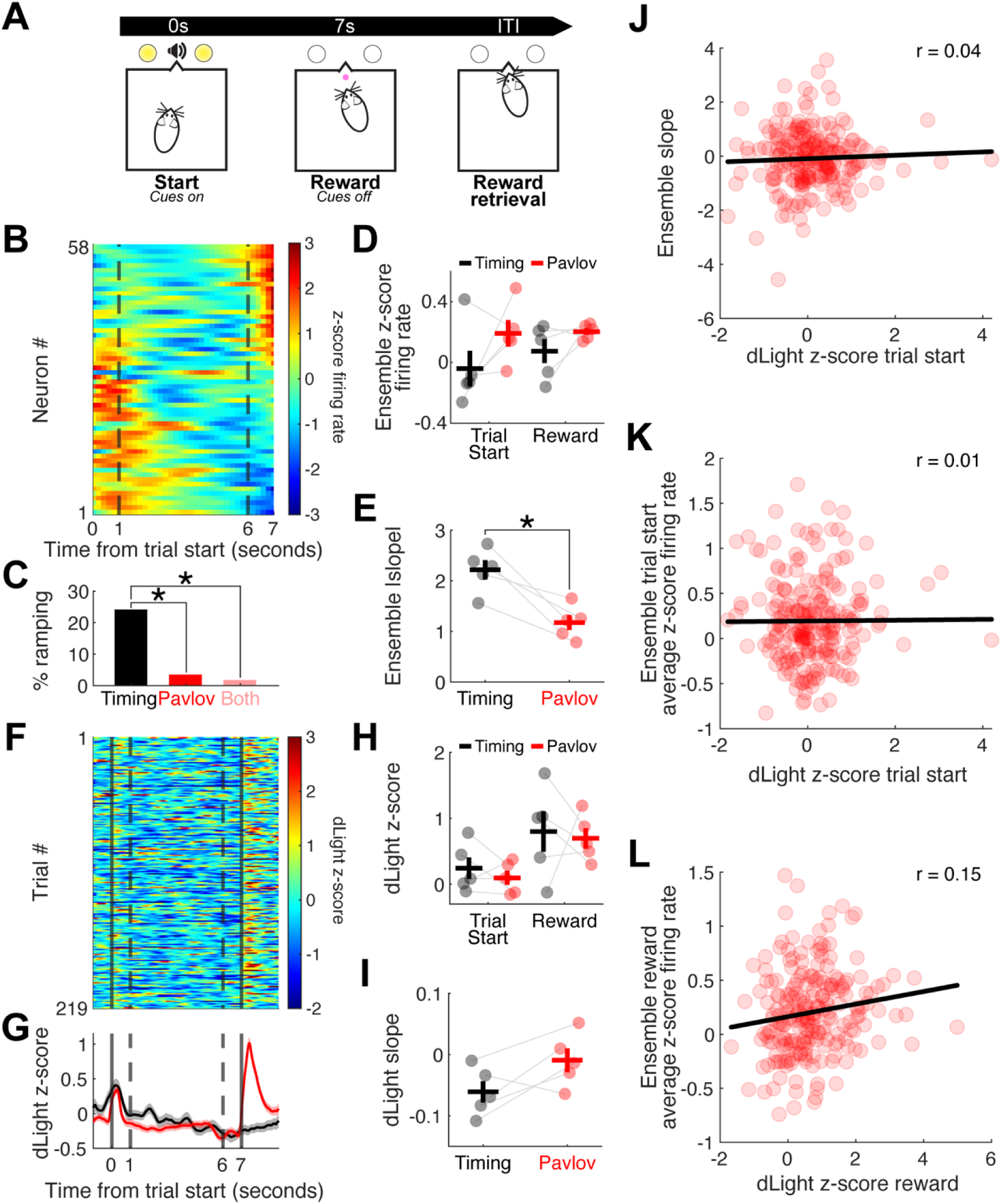
Dopamine and MSN modulations do not reflect anticipation of reward. **A)** Following recordings during the interval timing task, we recorded striatal dopamine (dLight z-score) and MSNs during a Pavlovian conditioning task. Cues randomly turned on for 7 seconds, and a sucrose pellet was delivered at cue offset. **B)** Peri-event time histogram of Pavlovian conditioning trials. MSNs are sorted by principal component 1. Dashed lines indicate the time interval for slope calculations in E and J. **C)** Fewer MSNs ramped during Pavlovian conditioning trials than during interval timing switch trials. Data from 58 MSNs in 5 mice. **D)** The average ensemble firing rate (z-score) in the second following trial start and reward delivery during interval timing switch trials (black) and Pavlovian conditioning trials (red). **E)** The average |slope| of all MSNs in an ensemble was higher during interval timing switch trials (black) than during Pavlovian conditioning trials (red). **F)** dLight z-score during every Pavlovian conditioning trial. Trials are aligned to trial start. Dashed lines indicate the time interval (1 – 6 seconds) for slope calculations in (I). **G)** Average dLight signal across switch trials during the interval timing task (black) and across Pavlovian conditioning trials (red). Dashed lines indicate the time interval for slope calculations in (I). Shading represents s.e.m. **H)** The average dLight z-score in the second following trial start and reward delivery in interval timing switch trials (black) and Pavlovian conditioning trials (red). **I)** The slope of the dLight signal from 1 to 6 seconds during interval timing switch trials (black) and Pavlovian conditioning trials (red). Dots represent individual mice, horizontal lines represent group averages, and vertical lines represent s.e.m. in D, E, H, and I. **J)** The average dLight z-score at trial start did not predict MSN ensemble slope during Pavlovian conditioning. Data from 58 MSNs in 5 mice. **K)** The average dLight z-score at trial start did not predict the average trial start firing rate of MSN ensembles. Data from 58 MSNs in 5 mice. **L)** The average dLight z-score at reward delivery was predictive of the reward delivery MSN ensemble firing rate. Data from 58 MSNs in 5 mice. Dots represent individual trials in J-L. Data in all panels from 98 Interval timing trials and 219 Pavlovian trials in 5 mice. \**p* < 0.05

Striatal MSNs were modulated around trial start or reward delivery. Fascinatingly, few neurons showed ramping activity during the interval (Fig 5B-C; timing vs. Pavlovian: *p* = 0.001; timing vs. both: *p* = 0.0003), as in our prior work^22^. Average ensemble firing rate at trial start and reward did not differ significantly between interval timing and Pavlovian conditioning trials (Fig 5D; trial start: *p* = 0.22, reward: *p* = 0.19). However, average MSN ensemble |slope| from 1 to 6 seconds was significantly lower during Pavlovian conditioning compared to interval timing (Fig 5E; *p* = 0.005; first second omitted to remove cue-related modulation).

Similarly, striatal dopamine, as measured by dLight z-score, showed strong phasic modulation at trial start and reward delivery during Pavlovian conditioning (Fig 5F-G). These modulations did not differ from those during interval timing trials (Fig 5H; trial start: *p* = 0.44, reward: *p* = 0.80). dLight signal during Pavlovian conditioning trials showed relatively little modulation between trial start and reward and the slope of the signal from 1 to 6 seconds was nearly zero (Fig 5I; average slope: -0.009 ± 0.04), which trended to be closer to 0 than the dLight slope from 1 to 6 seconds during interval timing switch trials (average slope: -0.06 ± 0.04; slope interval timing vs. Pavlovian: *p* = 0.08).

Finally, we asked if dLight z-score predicted MSN ensemble activity during Pavlovian conditioning. Unlike during interval timing, where dLight z-score early in the trial predicted ensemble slope (Fig 4I-J), trial start dLight modulations did not predict MSN ensemble slope from 1-6 seconds during Pavlovian conditioning (Fig 5J; r = 0.04, *p* = 0.51). dLight signal also did not predict MSN ensemble activity at trial start (Fig 5K; r = 0.01, *p* = 0.91), but reward dLight signal predicted MSN ensemble reward activity (Fig 5L; r = 0.15, *p* = 0.03). Together, these findings support past work^22^ showing that timing-related modulation during the interval timing task did not readily reflect anticipation of upcoming reward.

### SNc dopamine inhibition disrupts MSN ramping

Next, we wanted to causally prove the relationship between dopamine and MSN ramping activity. We inhibited dopamine neurons in the substantia nigra pars compacta (SNc) with Halorhodopsin during random switch trials, while recording MSN ensemble activity (Fig 6A-B). Inhibiting SNc dopamine neurons delayed switch response times (Fig 6C, Extended Data Fig. 6A, Extended Data Fig. 7A-B; laser off vs. laser on: *p* = 0.02), but did not affect movement trajectories, movement velocity (0 – 18 seconds: *p* = 0.23; 0 – 6 seconds: *p* = 0.27), or nosepokes (Fig 6D-F, Extended Data Fig. 5). This finding is consistent with extensive work from our group and others demonstrating that disrupting nigrostriatal dopamine, pharmacologically blocking dopamine receptors, or optogenetically disrupting D1 or D2 MSNs slows interval timing, increasing response times^15,23,33–35^.

**Figure 6.**
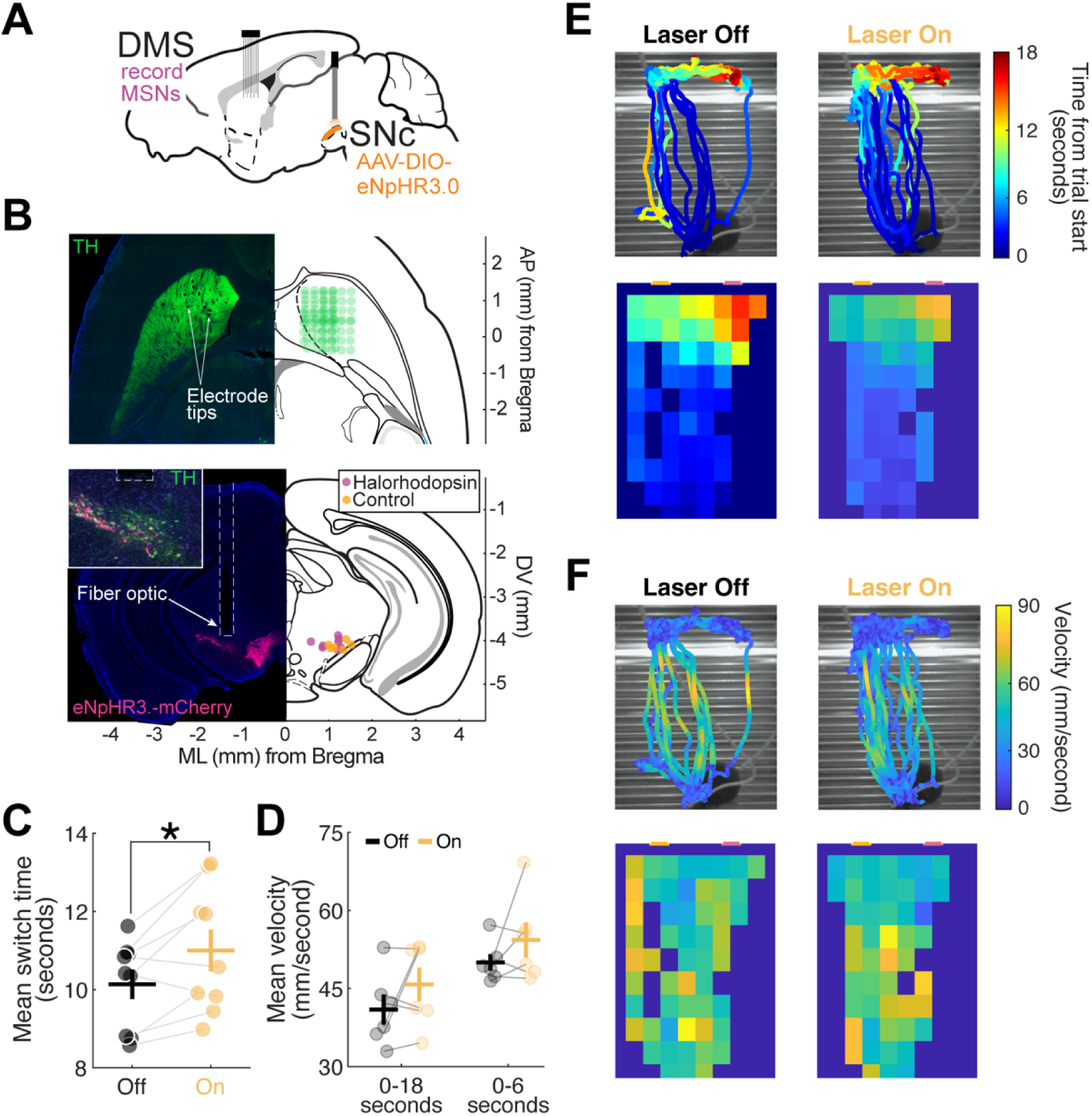
SNc dopamine inhibition disrupts timing but not movement. **A)** Top: We recorded MSNs in the DMS of DAT^IRES^-Cre mice while randomly inhibiting SNc dopamine neurons expressing Halorhodopsin (eNpHR3.0; 594 nm, 2 mW) from 0-18 seconds on 50% of trials. **B)** Top left: Horizontal section of DMS with electrode implant. Arrows identify electrode tracts. Top right: Horizontal section depicting electrode reconstruction from 6 mice with recording arrays implanted in the DMS. Bottom left: Coronal section of SNc with Halorhodopsin expression (pink). Dashed white box identifies fiber tract. Inset shows zoomed in section of SNc with merged stain of eNpHR3.0-mCherry (pink) and TH (green). Bottom right: Coronal section depicting approximate fiber optic tip locations above SNc. Pink circles represent locations from mice expressing Halorhodopsin, while yellow circles represent locations from control mice expressing no-opsin viruses. **C)** Optogenetic inhibition of dopamine neurons (2 mW) increased switch response times. 479 laser off trials and 341 laser on trials in 9 mice. **D)** Average movement velocity of mice did not differ between laser off (black) and laser on (yellow) trials when averaged across the full trial (0 – 18 seconds) and averaged over 0 to 6 seconds. Data from 115 laser off trials and 141 laser on trials in 6 mice. **E)** Top: DeepLabCut-tracked movement trajectories from a single session during laser off (left) and laser on (right) trials. Color represents time in trial (seconds). Bottom: Color represents the average time in trial (seconds) when mice were in each block for laser off (left) and laser on (right) trials. **F)** Top: DeepLabCut-tracked movement trajectories from a single session during laser off (left) and laser on (right) trials. Color represents velocity (mm/second). Bottom: Color represents the average velocity (mm/second) of mice within each block for laser off (left) and laser on (right) trials.

MSNs primarily ramped down with the laser off (Fig 7A-C; 70% down-ramping, 30% up-ramping), consistent with dLight-MSN recordings (see Fig. 1G). Intriguingly, we discovered that dopamine inhibition shifted this distribution, decreasing the percentage of down-ramping MSNs and increasing the percentage of up-ramping MSNs (Fig 7B-C; 49% down-ramping, 51% up-ramping: *p* = 0.01). However, further examination revealed that this shift was primarily driven by a reduction in the percentage of down-ramping MSNs exhibiting significant temporal modulation of firing rate, as defined by GLM analysis (Extended Data Fig. 7C-D). In contrast, dopamine inhibition did not affect the percentage of time-modulated up-ramping MSNs. Consistent with this selective effect, down-ramping MSN slopes shifted towards zero with dopamine inhibition, but up-ramping MSN slopes were unaffected (Fig 7D; down-ramping: *p* = 3.04×10^-5^; up-ramping: *p* = 0.26). This effect was absent in down-ramping MSNs of control mice expressing non-opsin virus in SNc dopamine neurons, although there were fewer recorded MSNs (Extended Data Fig. 6B-C). Together, these findings suggest that dopamine inhibition preferentially attenuates the monotonic decrease in firing rate of down-ramping MSNs, rather than simply reversing the direction of ramping activity.

**Figure 7.**
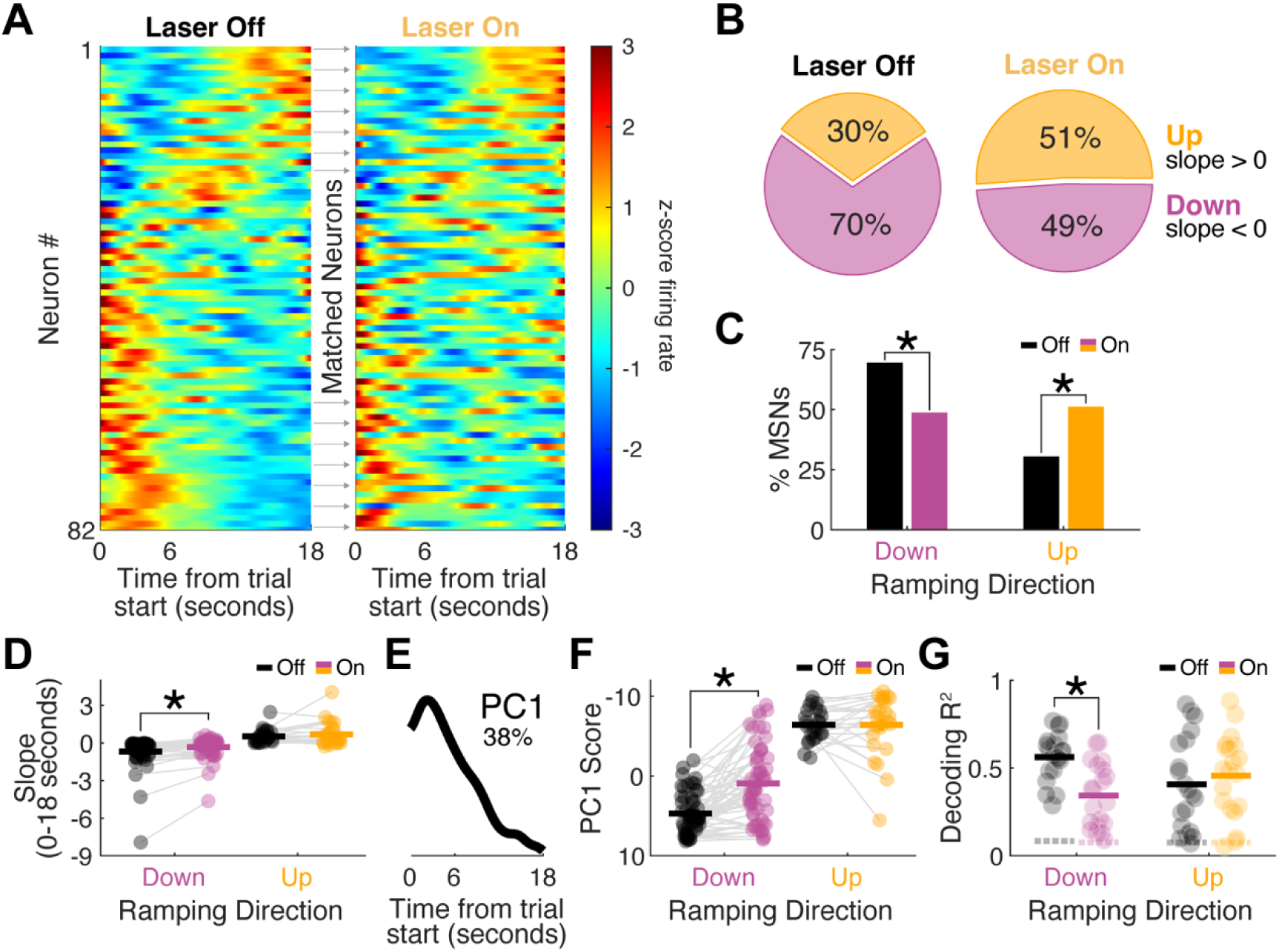
SNc dopamine inhibition disrupts MSN ramping. **A)** PETHs of interval timing switch trials with the laser off (left) and with the laser on (right). MSNs are sorted by PC1 for the laser off condition and displayed in the same order for the laser on condition. **B)** Percentage of MSNs that are up-ramping (yellow) and down-ramping (pink) with the laser off (left) and with the laser on (right). Values in pie charts are rounded to the nearest whole number. **C)** Laser inhibition decreased the percentage of down-ramping MSNs and increased the percentage of up-ramping MSNs. Data in A-C from 130 laser off trials and 111 laser on trials from 82 MSNs in 6 mice. **D)** The slope of 57 down-ramping MSNs increased with laser inhibition (pink), but slope did not change in 25 up-ramping MSNs (yellow). **E)** Principal component analysis (PCA) found that principal component 1 (PC1) showed ramping activity and explained 38% of the variance. **F)** PC1 scores in 57 down-ramping MSNs decreased with laser inhibition (pink) but did not change in 25 up-ramping MSNs. Y axis is flipped to match slope in (D). **G)** Down-ramping MSN ensembles during laser inhibition trials (pink) had weaker temporal decoding than during laser off trials. Temporal decoding in up-ramping MSN ensembles did not differ between laser off and laser on (yellow) trials. Dashed lines represent group averages from shuffled data. Data in all panels from 6 mice. \**p* < 0.05.

As a sensitivity analysis, we turned to PCA to characterize patterns of neuronal activity with dopamine inhibition. As reported previously^15,22,25,27,32^, PC1 showed ramping activity and explained 38% of the variance (Fig 7E). We found that PC1 scores in individual down-ramping MSNs and in down-ramping ensembles decreased with dopamine inhibition, but again, PC1 scores in individual up-ramping MSNs and ensembles were unaffected (Fig 7F, Extended Data Fig. 7E; down-ramping MSNs: *p* = 1.20×10^-8^; up-ramping MSNs: *p* = 1.00). Finally, we evaluated how well ensembles decoded time in laser off versus laser on trials (Fig 7G, Extended Data Figure 7F-G). Dopamine inhibition weakened temporal decoding in down-ramping MSNs (*p* = 0.0005) but did not affect temporal decoding in up-ramping MSNs (*p* = 0.58). Dopamine inhibition’s specific effects on down-ramping dynamics are consistent with prior findings that down-ramping predicts switch response time and that dopamine preferentially predicts down-ramping rather than up-ramping MSN ensembles. In summary MSN recordings, dopamine recordings, and dopamine inactivation provide convergent and complementary evidence that dopamine supports time-dependent ramping specifically in down-ramping MSNs.

### Perturbing drift-diffusion dynamics predicts timing in mice and humans with Parkinson’s disease

Neural activity integrates temporal evidence to guide timed responses^15,22,25,27,31^. In the striatum, this process can be captured by drift-diffusion, with D1-MSNs exhibiting downward drift rates, and D2-MSNs exhibiting upward drift rates^15^. We applied this model to our data with optogenetic inactivation of SNc dopamine, guided by the effects on down-ramping MSN activity.

We adopted a drift-diffusion model (DDM) described in detail previously^15^, where *x* represents the neuronal firing rate of an ‘output’ unit collecting evidence across striatal MSNs (Fig. 8A). In this model, *t* represents time measured in seconds, and *dx* and *dt* mean “change” in *x* and *t* respectively (equivalent to the derivative *dx*/*dt*):

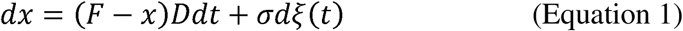

**Figure 8.**
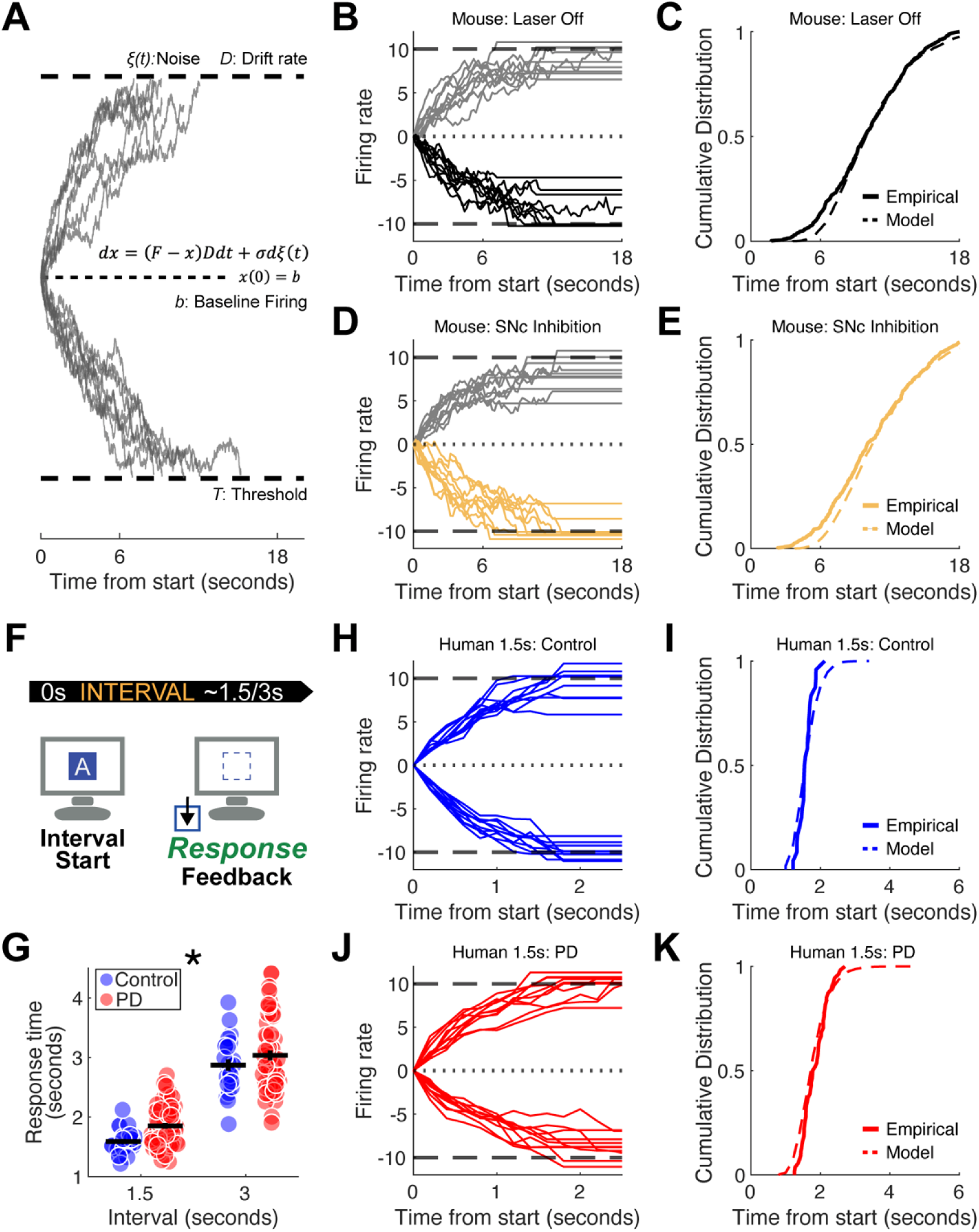
DDMs explain interval timing perturbations from mice to humans. **A)** Schematic representation of the drift-diffusion model (DDM)^15^ governed by the stochastic differential equation *dx* = (*F – x*)*Ddt* + *σdξ*(*t*). Up-ramping and down-ramping processes accumulate evidence over time until reaching a decision threshold (*T*). **B)** Simulated DDM trajectories and **C)** corresponding cumulative distribution function (CDF) fits against empirical switch times for mice during baseline performance (black; same data as optogenetics laser off in Fig 6C). **D)** DDM trajectories and **E)** CDF fits during optogenetic inhibition of SNc dopamine, accurately modeled solely by an increase in the down-ramping drift rate parameter (*D_down_*). **F)** Parkinson’s disease patients and control participants performed an interval timing task in which they indicate their estimation of elapsed time with a keypress response. Following a variable inter-trial interval, instructions appeared on a computer screen indicating whether patients should time a short (1.5 second) or long (3 second) interval. Subsequently, a blue square appeared to indicate interval start. Patients responded with a keypress when they estimated the target interval had elapsed, upon which the blue box would disappear. In one third of trials, patients received feedback on the accuracy of their response. **G)** Human interval timing behavior demonstrating mean response times for 24 healthy control participants (blue) and 54 Parkinson’s disease participants (red) at 1.5-second and 3.0-second intervals. As with SNc dopamine inhibition, PD patients are slower across both intervals. **H)** DDM trajectories and **I)** CDF fits for healthy control participants timing the 1.5-second interval. **J)** DDM trajectories and **K)** CDF fits for PD patients timing the 1.5-second interval. Mirroring the mechanistic model in mice, the temporal deficits observed in PD patients are successfully captured entirely by a selective alteration to the downward drift rate. * *p* < 0.05

The model has three independent parameters *F, D*, and *σ*, and a threshold value *T* defined by

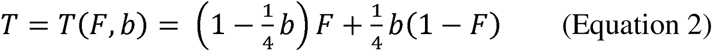

The baseline firing rate *b* is set initially at 0 then driven by input *F* similar to the dendritic current induced by the overlap of corticostriatal postsynaptic potentials^38^. With each unit of time *dt*, we suppose there is corticostriatal stimulation that provides incremental input proportional to the activity itself, (*F* – *x*) *D,* to reach a decision threshold. *D* is positive or negative *drift rate* for the event sequence and is inversely proportional to each neuron’s integration time constant. The drift, together with noise *ξ*(*t*) (of zero mean and strength *σ*), leads to fluctuating accumulation that eventually crosses a decision threshold *T* (see Equation 2). The time *t*\* elapsed for the firing rate to reach the threshold, *x*(*t*\*) = *T* models the *switch response time*.

As in past work, drift-diffusion models (DDMs) accurately captured empirical behavior during baseline performance (*R*^2^ = 0.99; Fig 8B-C). Extraordinarily, we discovered that changing a single parameter – drift rate (*D_down_*) by 8% – could accurately model behavioral deficits with optogenetic inhibition (from -0.21 to -0.23; *R*^2^ = 0.98; Fig 8D-E). All other model parameters—including noise (1), thresholds (±10), input drive (±2), and up-ramping drift (0.21)—were held constant. These fascinating data suggest that small, unidirectional changes in drift rate could replicate the effects of dopaminergic inhibition.

To determine if DDMs were relevant to human behavior, we examined interval timing in human Parkinson’s disease (PD). We recruited 24 human control participants and 54 PD participants to perform an interval timing task with 1.5 or 3 second intervals as in our prior work (Fig 8F)^39,40^. A linear mixed-effects model to evaluate these response times revealed a significant main effect of both disease (*F*(_1,152_) = 7.43, *p* = 0.007) and interval (*F*(_1,152_) = 284.49, *p* < 0.001). Crucially, there was no significant disease-by-interval interaction (*F*(_1,152_) = 0.41, *p* = 0.51). Model fit criteria favored the additive fixed-effects model (AIC = 201) over the interaction model (AIC = 203), demonstrating that the magnitude of PD-related slowing is consistent across both the 1.5 second and 3.0 second intervals (Fig 8G). Age differed between PD and control participants (*F*(_1,152_) = 7.20, *p* = 0.008); however, the main effect trended when age was added to our model (*F*(_1,152_) = 3.57, *p* = 0.06). Sex and years of education did not differ between groups and, accordingly, main effects of disease and interval remained significant with these included in models.

As in mice, DDMs predicted human response times in both the 1.5 (Control *R*^2^ = 0.89; PD *R*^2^ = 0.93) and 3.0 second intervals (Control *R*^2^ = 0.95; PD *R*^2^ = 0.97) (Fig. 8H-I and Extended Data Fig. 8A-B). Mirroring the mechanistic modeling in mice, the temporal deficits observed in PD patients were captured entirely by selectively altering the downward drift rate. For the 1.5-second interval, *D_down_* increased by 25% (from 0.80 in controls to 1.00 in PD patients; Fig. 8J-K). For the 3.0-second interval, *D_down_* similarly increased by 4% (0.20 in controls to 0.21 in PD patients; Extended Data Fig. 8C-D). As with mice, other parameters such as noise (1.5), thresholds (±10), input drive (±10), and up-ramping drift (0.8) were held constant. Taken together, these results suggest that dopaminergic dysfunction alters specific trajectories of cognitive computations, and that shifts in down-ramping drift robustly predict timing deficits across species.

## DISCUSSION

We investigated the role of dopamine in cognition by recording dopamine dynamics via dLight fiber photometry during interval timing. We tested the hypothesis that dopamine affects MSN ramping activity that predicts temporal control of action. Consistent with past work, we found that MSNs encoded time, but only down-ramping MSNs predicted animals’ internal estimates of time. We found that dopamine dynamics, as measured by dLight, did not robustly predict time but did predict timed response behavior as well as down-ramping MSN ensemble activity. Furthermore, we discovered that inactivating dopamine primarily affected down-ramping MSN dynamics. These data could be accounted for by changes in downward drift-diffusion dynamics, which explained both interval timing behavior in mice and in humans with Parkinson’s disease. Together, these data illustrate how dopamine affects drift-diffusion dynamics in the striatum. Indeed, by decreasing down-ramping dynamics while preserving up-ramping dynamics, we find that the aggregate vector of ensemble MSN activity is shifted by disrupting dopamine. These data illustrate how striatal dopamine can instantiate cognitive processing and could be helpful in further understanding striatal circuits towards development of new diagnostic or therapeutic approaches for human disease.

Past work by our group and others has implicated that MSNs expressing D1-dopamine receptors ramp down, while D2-dopamine-receptor expressing MSNs ramp up^15,41^. Here, we observe that phasic dopamine as measured by dLight z-score ramps down towards timed responses, that dopamine is correlated with down-ramping MSNs, and that dopamine inactivation primarily affects down-ramping MSNs. Furthermore, DDMs suggest that perturbations in down-ramping drift rates are sufficient to explain behavioral changes in mice with dopamine disruptions and in human PD patients who have disrupted nigrostriatal dopamine. One model for these data is that dopamine exerts excitatory modulatory influence on D1-MSNs, as our data suggest that dopamine acts primarily on down-ramping, or putative D1-MSNs. D2-MSNs may play myriad other roles; and we note that up-ramping, or putative D2-MSNs, still represent elapsed time. D2-MSNs and D1-MSNs are often co-activated during movement^42,43^. Further disambiguating dopamine vs. D1-MSN and D2-MSN activity requires complex intersectional and optogenetic strategies^44^; our results motivate these efforts and suggest that dopamine primarily affects D1-MSN ramping during interval timing.

Ramping is ubiquitous in corticostriatal circuits^13,22,24,27,41,45,46^. While striatal ramping requires cortical input^21,32^, corticostriatal projections tend to ramp up and are excitatory^47^. These may preferentially influence D2-MSNs, which would ramp up as excited by corticostriatal projections. Additionally, D2-MSNs can laterally inhibit D1-MSNs^48,49^, which may exert response control via down-ramping D1-MSNs. Notably, our strongest results are at the ensemble rather than single neuron level, which suggests that striatal ensembles may integrate large volumes of corticostriatal input to drive responses. Recording from and testing the relative roles of D1- vs. D2-MSNs again requires complex intersectional optogenetics but may be required to unravel striatal circuit contributions to interval timing.

An advance of our work is to record dopamine in conjunction with MSN activity. Our dopamine signals trended down over the interval, contrary to other studies that report up-ramping dopamine dynamics in advance of reward^50,51^. Notably, our view does not support canonical contributions of dopamine to firing rate or complementary effects of D1-MSNs vs D2-MSNs^52–55^. Rather, our data suggest that dopamine specifically shifts downward ramping trajectories. Notably, dLight in the DMS was related to movement velocity but could not entirely explain the effects on ramping dynamics. These data highlight a specific contribution of dopamine to interval timing. DDMs can explain a broad range of cognitive behaviors including decision making and working memory, but it remains to be seen how dopamine in the striatum affects these dynamics.

Our work has several limitations. First, we conducted dLight fiber photometry in DAT^IRES^-Cre mice, and we could not isolate D1- or D2-MSNs as noted above. Second, dLight only measures relative dopamine signals, although we note that optogenetic inhibition did reliably affect dLight activity. Third, a larger patient cohort would be necessary to fully account for contributions of age, dopaminergic medication, and other clinical factors to our findings. Finally, while we translate our work from mice to humans with PD, it is unclear if the DDM dynamics reported here are relevant for other cognitive operations.

In summary, we demonstrate that dopamine specifically shifts downward drift-diffusion dynamics during interval timing in mice and can account for impaired interval timing in human PD. These data provide insight into how dopamine affects cognitive processing in the striatum, which is highly relevant for a better understanding of dopaminergic diseases affecting cognition.

## MATERIALS AND METHODS

### Mice

All procedures were approved by the Institutional Animal Care and Use Committee (IACUC) at the University of Iowa (#3052039). A total of 34 mice (10 excluded) were used across all mouse experiments (see Supplementary Table 1). Mice had access to *ad libitum* water, were food restricted to ∼85% of their starting bodyweight, and were housed individually post-surgery in accordance with methods described in detail previously^15,22,24^. Four mice were excluded from the combined fiber photometry/electrophysiology experiment due to low dLight signal-to-noise (*n* = 1), inconsistent electrophysiology recording (*n* = 1), or insufficient behavior during recordings (*n* = 2). Four mice were excluded from the combined optogenetics/electrophysiology experiment, and two mice were excluded from the optogenetics only experiment due to lack of opsin expression or mistargeting of the fiber optic. Electrophysiology data from separate recording sessions in 8 mice were published previously^22^.

### Interval timing switch task

We used a mouse-optimized interval timing task described in detail previously^15,22–24,56^. Briefly, mice were trained in sound-attenuating chambers with a nosepoke on the back wall, and two additional nosepokes flanking a central food magazine on the front wall (Fig. 1A; MedAssociates, St. Albans, VT). Cue lights were positioned above the two front nosepokes, and all nosepokes contained internal lights and infrared beams to register entries and exits. Mice began training with fixed-ratio nosepoke response trials to earn rewards (20 mg sucrose pellets; Bio-Serv, Flemington, NJ).

Following fixed-ratio training, mice began training in the interval timing switch task. Following a variable inter-trial interval (ITI), mice initiated trials by responding at the back nosepoke, which triggered cue lights and an auditory tone (3kHz, 72 dB). Cues were identical on all trials and sustained for the entire trial duration. On 50% of trials (randomly assigned), cues sustained for 6 seconds and mice were rewarded for the first response at the designated “short” nosepoke after 6 seconds. On the other 50% of trials cues sustained for 18 seconds and mice were rewarded for the first response at the designated “long” nosepoke after 18 seconds after at least one short nosepoke. During these *switch trials*, mice first responded at the “short” nosepoke before *switching* to the “long” nosepoke. The time the mouse exited the “short” nosepoke before moving to the “long” nosepoke is defined as the *switch response time*. For all analyses, only correct switch trials were included.

### Pavlovian conditioning task

After interval timing recordings, mice were trained in a Pavlovian conditioning task as described in detail previously^22^. Briefly, during 30-minute sessions mice were randomly presented with auditory (6 kHz, 72 dB) and light cues for 7 seconds. At cue termination, mice were delivered a sucrose pellet reward. There was an average of 9.4 ± 1.1 calendar days between the interval timing recording and Pavlovian recording. The *average reward rate* (rewards/minute) and total *number of rewards* earned per session were recorded.

### Movement tracking

Behavioral sessions were recorded at 30 or 60 frames per second with OBS Studio software (version 27.1.3) with a USB camera (Arducam, Nanjing, China) positioned above the behavioral chamber. Videos were analyzed with DeepLabCut (version 2.2rc3) to track 2-D position of mice during trials^26^. As detailed in prior work^22^, these data were processed using custom MATLAB code to remove and correct invalid points using interpolation via piecewise cubit Hermite polynomials (*pchip*) to produce smooth trajectories. Positional data in pixels was converted to real-world points (millimeters) using video calibration with a 1 x 1-cm squares checkerboard and built-in MATLAB functions (*detectCheckerboardPoints, estimateExtrinsics, img2world2d*). *Movement velocity* was calculated as the Euclidean distance in consecutive frames divided by the inter-frame interval (1/30 or 1/60 seconds).

### Stereotaxic surgery

Surgical methods were identical to those in previous reports^15,22,24^. Mice were anesthetized with inhaled 4% isoflurane and maintained at 1-2% isoflurane during surgery. Depending on the experiment, craniotomies were drilled above the dorsomedial striatum (DMS) and/or substantia nigra pars compacta (SNc), according to the coordinates stated below. Viruses were infused at a rate of 0.05 µl/min (Legato 130 Syringe Pump, KD Scientific, Holliston, MA). Skull screws were used to anchor the surgical headcap for all implant surgeries. Implants and screws were sealed with Slo-Zap cyanoacrylate accelerated with Zip-Kicker (Pacer Technologies, Rancho Cucamonga, CA) and methyl methacrylate (AM Systems, Port Angeles, WA). Mice were given at minimum 1 week for postoperative recovery before beginning food restriction and behavioral training.

For fiber photometry with electrophysiology experiments, a craniotomy was drilled above the DMS (AP +0.5, ML ±1.4; 4 mice in the right hemisphere and 4 mice in the left hemisphere). AAV5-CAG-dLight1.3b was injected (0.25 μl each; DV -2.8, DV -2.7) before implanting a multi-channel electrode with a central fiber optic (fiber photometry “optrode”; DV -2.7; Microprobes, Gaithersburg, MD). The electrode was grounded to at least one skull screw.

For optogenetics, AAV5-Ef1a-DIO-eNpHR3.0-mcherry (Halo) or AAV5-Ef1a-DIO-mcherry (control) was bilaterally injected into the SNc (AP -3.0, ML ±1.3, DV -4.5, DV -4.25). Fiber optics were implanted bilaterally into the SNc (AP -3.0, ML ±1.3, DV -4.3). In 8 mice, craniotomies were also made above the DMS as described above to implant multi-channel electrodes. In 2 mice, Halo was injected only unilaterally in the SNc as above, AAV5-CAG-dLight1.3b was injected unilaterally into the DMS (ipsilateral to the SNc injection hemisphere; AP +0.5, ML ±1.4, DV -2.7 DV -2.6), and fiber optics were implanted above SNc (DV -4.3) and DMS (DV -2.6).

### Fiber photometry

Mice were tethered to a unilateral optical patch cord connected to a fiber photometry system (Doric Lenses Inc, Quebec, Canada) with two excitation wavelengths (405-nm LED isosbestic signal modulation at 208.616 Hz and 470-nm LED dLight signal modulation at 572.205 Hz). Isosbestic and dLight fluorescence wavelengths were measured via integrated LED mini cubes and photo detector (iLFMC4, Doric Lenses Inc). Recording sessions lasted 90 minutes. Changes in dLight signal were downsampled to 30 Hz, low-pass filtered at 5 Hz, detrended to compensate for exponential decay and photobleaching, motion corrected using the isosbestic signal, z-scored to compare across mice, and aligned to task events.

To test the effect of optogenetic dopamine neuron inhibition with Halorhodospin on DMS dopamine release, we randomly turned on a 594 nm (2 mW) laser for 2 or 7 seconds while recording dLight. To compare dLight signal during laser off and laser on conditions we calculated the area under the curve (*trapz*) during 2 second or 7 second laser on trials and for 2 or 7 second periods of laser off.

In interval timing recordings, we calculated the average dLight signal, the slope of the dLight signal (*polyfit*), and Δ velocity in the four second period preceding switch responses for each trial. We also calculated the slope of the dLight signal in the four second period preceding short nosepoke releases that were not switch responses. For correlations (*corrcoef*) of dLight signal, dLight slope, and Δ velocity with switch response time or velocity, we excluded trials with switch response times before 4 seconds and used Cook’s distance to remove outliers (*fitlm*, threshold of 3*mean Cook’s D). To determine how velocity contributed to the relationship between dLight slope and switch response time, we ran partial correlations of dLight slope with switch time controlling for velocity (*partialcorr*) and used a Steiger’s z-test to compare the normal correlation to the partial correlation.

We also calculated the average dLight signal in the period before the switch response (pre-switch), the period after the switch response (post-switch), from 0 to 6 seconds, and from 6 to 12 seconds. We correlated the pre-six and pre-switch signals with MSN ensemble slope (see below) with *corrcoef*. For Pavlovian conditioning, we calculated the average dLight z-score in the 1 second periods following trial start and reward delivery. We also calculated and compared the slopes (*polyfit*) of dLight from 1 to 6 seconds during Pavlovian conditioning and interval timing trials.

### Neural recordings

We recorded single DMS MSNs using a multi-electrode recording system (Supplementary Table 2). Recordings were processed using Plexon Offline Sorter (version 4.6.0, Plexon, Dallas TX) to remove artifacts and single neurons were identified using PCA and waveform shape. We used the following criteria to identify single neurons: 1) consistent waveform shape; 2) a separable cluster in PCA space; and 3) a consistent refractory period of at least 2 milliseconds in interspike interval histograms. Putative medium spiny neurons were separated from fast-spiking interneurons based on hierarchical clustering of the waveform peak-to-trough ratio and the half-peak width (*fitgmdist* and *cluster*)^15,22,27,32^. As described previously^22,24^, we sorted the same neurons across two separate interval timing sessions or across separate interval timing and Pavlovian conditioning sessions for each mouse. To verify sorting integrity across the two sessions, we quantitatively compared waveform similarity via correlation coefficients and removed neurons with correlation coefficients below 0.8 (data for 22/130 neurons removed for two interval timing task sessions; data for 2/60 neurons removed for interval timing and Pavlovian task sessions). Only medium spiny neurons were analyzed.

We harnessed trial-by-trial generalized linear models (GLMs; *fitglme*) to identify modulation patterns of individual neurons. For each trial, we calculated kernel density estimates of firing rates across the interval (0 to 18 seconds for interval timing or 0 to 7 seconds for Pavlovian conditioning) binned at 0.2 seconds, with a bandwidth of 0.2. We classified MSNs based on the sign of their GLM-derived slopes from 0 to 18 seconds: neurons with negative slopes were classified as ‘down-ramping’, whereas neurons with positive slopes were classified as ‘up-ramping’ (independent of whether the GLM-derived slope reached statistical significance). As a sensitivity analysis, we focused on MSNs with significant linear fits from 0 to 18 seconds; although this decreased statistical power for comparisons, particularly of up-ramping MSNs. We used this definition to compare the percentage of ramping neurons in interval timing vs. Pavlovian conditioning sessions (Fig. 5) and for the analyses presented in Extended Data Fig. 7C-D.

To efficiently fit individual trial-level data, slope of firing rate (0 to 18 seconds, 0 to 6 seconds, or 1 to 6 seconds) was calculated using polynomial curve fitting (*polyfit*). To calculate ensemble slopes, we averaged individual slopes across all neurons from a single mouse for each trial. To calculate firing rate at trial start and reward delivery, we averaged the z-scored estimated firing rate during the 1 second period after trial start or reward delivery for each trial. Correlations between MSN slope/firing rate and switch time/dLight z-score were performed at the trial level, with each data point representing the slope/firing rate and corresponding switch time/dLight z-score from an individual trial. Thus, MSNs had multiple observations, with the number of observations corresponding to the number of trials recorded for each neuron.

To examine moment-by-moment correlations between firing rate and dLight signal, we binned firing rate at 0.2 seconds from -6 seconds before trial start to 24 seconds after trial start and calculated an estimated firing rate using a 0.3 second bandwidth to match dLight signal sampling rate at 30 Hz. We then calculated Pearson’s correlation coefficients (*corrcoef*) between dLight signal and firing rate using a sliding window calculation, with a window size of 3 seconds and a step of 0.5 seconds.

Finally, we used principal component analysis (PCA) to identify data-driven patterns of z-score neuronal activity and focused on principal component 1 (PC1), which explained the most variance. As in previous work^15,22,25,27,32^, PC1 displayed time-dependent ramping activity.

### Temporal decoding

We constructed naïve Bayesian classifiers to evaluate how neuronal ensembles and dLight signal predicted time^10,15,22,25,27,32^, which predicted objective time from firing rate or dLight z-score during a trial using leave-one-out cross validation. We ran the decoding analysis on different neuronal ensembles based on ramping direction (all neurons, down-ramping neurons, or up-ramping neurons). To correct for ensembles of different sizes, we ran the classifier on 200 iterations of an equivalent number of randomly selected neurons from each group (30 MSNs/group for comparing all MSNs, down-ramping MSNs, and up-ramping MSNs; 6 MSNs/group for comparing neuronal ensembles to dLight; and 20 MSNs/group for optogenetics). For dLight, we binned the z-scored signal into 0.2 second bins to match neuronal firing rate. We quantified classifier performance by computing the R^2^ of objective time vs. predicted time for each trial. We also compared classifier performance to neuronal ensembles with time-shuffled firing rates or dLight signal with time-shuffled signal.

### Optogenetics

Mice were well-trained in the interval timing switch task before beginning optogenetic procedures. Mice were tethered to a 594-nm laser (Cobolt Mambo^™^ 100, HÜBNER Photonics, Kassel, Germany) controlled by a laser shutter control (SR470, Stanford Research Systems, Sunnyvale, CA) through a bilateral or unilateral optical rotary joint and optical patch cords (Doric Lenses Inc, Quebec City, Quebec). Laser light was randomly delivered during 50% of trials. Laser light was calibrated to either 2 mW or 5 mW at the fiber tip and was delivered for either 2 seconds at trial start or for the entire trial (0-6 or 0-18 seconds) in separate behavioral sessions. Sessions with 5 mW full trial inhibition were removed from analysis because of clear motor effects identified using DeepLabCut movement tracking.

### Human participants

A total of 54 participants with Parkinson’s disease (aged 48 to 87 years; 19 females) and 24 control participants (aged 40 to 78 years; 15 females) performed an interval timing task. All participants were recruited under a study protocol approved by the University of Iowa Institutional Review Board (IRB #202302747) and provided informed consent prior to participation. See Supplementary Table 3 for further demographic information.

### Human interval timing task

Participants completed a modified version of an interval timing task^40^ programmed in MATLAB using Psychtoolbox and presented on a Dell XPS workstation with a 19-inch monitor. The task required estimation of either a short (1.5 seconds) or a long (3 seconds) interval and participants were instructed to estimate these intervals without counting. The task consisted of three phases. During the demonstration phase, participants were instructed to simply observe as a blue box appeared and changed to a red box when the indicated interval (long or short) had elapsed. During the subsequent training and testing phases, trials began with instructional text informing participants to time the 1.5 second or 3 second interval. Following a 1-1.5 second delay, the beginning of the interval was marked by the appearance of a blue square, and participants were asked to press the keyboard spacebar when they judged the interval to have elapsed (Figure 8F). During each trial, the letter “A” was superimposed on the blue square at random intervals to distract from counting. Intertrial intervals were jittered (2-3 seconds), along with intervals between short/long cue text presentation and timed-stimulus onset (0.5-1.5 seconds). In the training phase, feedback was given on accuracy for every trial. During the testing phase, feedback was only given on a third of trials. Participants performed four blocks of 20 randomly interleaved short and long trials, with a break after each block.

### Drift diffusion model

Our DDMs were described in detail previously^15^ and simulate switch response times in mice and timing responses in human. DDMs were built in MATLAB, starting from initial value *b* and with discrete computer simulation steps 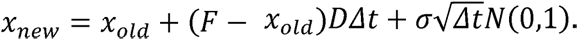 N(0,1) represents random values generated from the standardized Gaussian distribution (mean = 0 and standard deviation = 1). The integration timestep was taken *Δt =* 0.2 to be like the 0.2-second bin size used in the analysis of firing rate data. For each numerical simulation, the “switch response time” was defined as the time *t\** when variable *x* first reached the threshold value *T* = *F*(1 – *b*/4) + (1 – *F*)*b*/4 For each condition, we ran 3000 simulations of the model (up to 25 seconds per trial) and recorded the switch response times.

### Immunohistochemistry

Mice were transcardially perfused with 1X phosphate-buffered saline (PBS) and 4% paraformaldehyde (PFA) after anesthesia using ketamine (100 mg/kg IP) and xylazine (10 mg/kg IP). Before cryosectioning, brains were fixed in 4% PFA followed by 30% sucrose in PBS. Brains implanted with optrodes for fiber photometry/electrophysiology recordings were sliced horizontally. Brains for optogenetics with electrophysiology recordings were first cut in half and sections containing substantia nigra were sliced coronally while sections containing striatum were sliced horizontally. Striatal sections were stained for tyrosine hydroxylase and GFP with primary antibodies at 4°C (rabbit anti-TH, Millipore AB152, 1:1000, 24 hours or chicken anti-TH, Abcam ab76442, 1:1000, 48 hours; chicken anti-GFP, Abcam ab13970, 1:500, 24 hours or rabbit anti-GFP, enQuirebio, QAB10298, 48 hours) and visualized with Alexa Fluor fluorescent secondary antibodies at room temperature for 2 hours (goat anti-rabbit IgG Alexa 488, Invitrogen A11008, 1:1000 or goat anti-chicken IgG Alexa 405, Invitrogen A48260, 1:250; goat anti-chicken IgG Alexa 488, Invitrogen A11039, 1:1000 or goat anti-rabbit IgG 488, Invitrogen A11008, 1:1000). Midbrain sections were stained for tyrosine hydroxylase (rabbit anti-TH, Millipore AB152, 1:1000, 24 hours) and visualized with Alexa Fluor fluorescent secondary antibodies at room temperature for 2 hours (goat anti-rabbit IgG Alexa 488, Invitrogen A11008, 1:1000). Halorhodospin was imaged for native expression. To histologically confirm the location of electrode arrays, fiber implants, and virus expression, slices were imaged with VS-ASW-S6 imaging software (Olympus, Center Valley, PA). For one optrode mouse, only electrode coordinates, but not fiber optic coordinates, could be reconstructed from histological images.

### Statistics

All data and code will be publicly available via Zenodo upon publication. All statistical results are reported in Supplemental Table 4. We described data using group averages and standard error of the mean (s.e.m.). To compare dLight signal before and after 6 seconds or switch response, dLight or firing rate activity between interval timing and Pavlovian conditioning, and differences between optogenetics laser off and on conditions, we used paired t-tests (*ttest* in MATLAB). To compare R^2^ values from Bayesian classifiers, we fit linear models in R (*lmer*), running ANOVA to find main effects of neuron group, and making post hoc comparisons between groups using *emmeans*. To determine if correlations between moment-by-moment dLight and MSN firing rate were different than 0 we ran linear mixed effect models (*fitlme* in MATLAB) with post hoc contrast tests (*coefTest* in MATLAB). For comparisons of percentage of ramping neurons, we used Chi-squared tests (*crosstab* in MATLAB). For individual trial correlations, we used Pearson’s linear correlations (*corrcoef* in MATLAB). To determine if moment-by-moment dLight and velocity correlations were significantly different than 0, we used Wilcoxon signed rank tests (*signrank* in MATLAB). To compare correlations to partial correlations, we used Steiger’s z-tests (custom code in MATLAB). Finally, we evaluated human interval timing behavior by comparing linear mixed-effects models with 1) main effects of disease condition (control vs. PD), interval duration (1.5 vs. 3 seconds), sex, age, and years of education or 2) an interaction between disease condition and interval duration (*fitglme* in MATLAB).

