## Supplemental Figures for "Dopamine shifts striatal ramping dynamics"

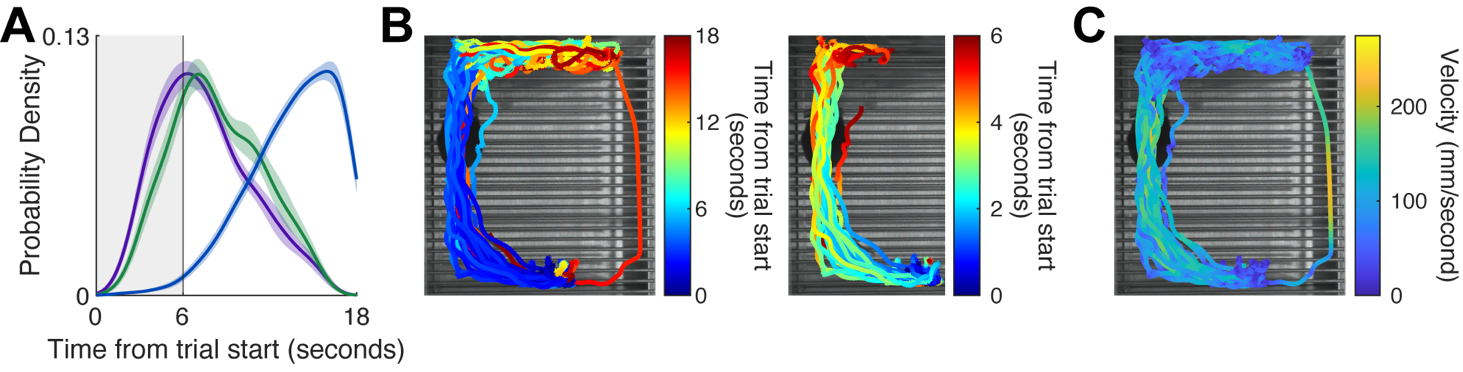

**Extended Data Figure 1.**

A) Response probability distribution for short nosepokes (purple), long nosepokes (blue), and switch responses (green). Data from 494 trials in 25 mice (13 female).
B) We tracked mouse position during trials. Colored lines represent the movement trajectories of a single mouse during all interval timing switch trials from a single session. Color represents the time within the trial aligned to trial start (seconds). Left: Movement trajectories across the full trial (0 to 18 seconds). Right: Movement trajectory during the first 6 seconds of the same trials.
C) We tracked mouse movement velocity during trials. Color represents velocity (mm/second) during interval timing switch trials from a single session.

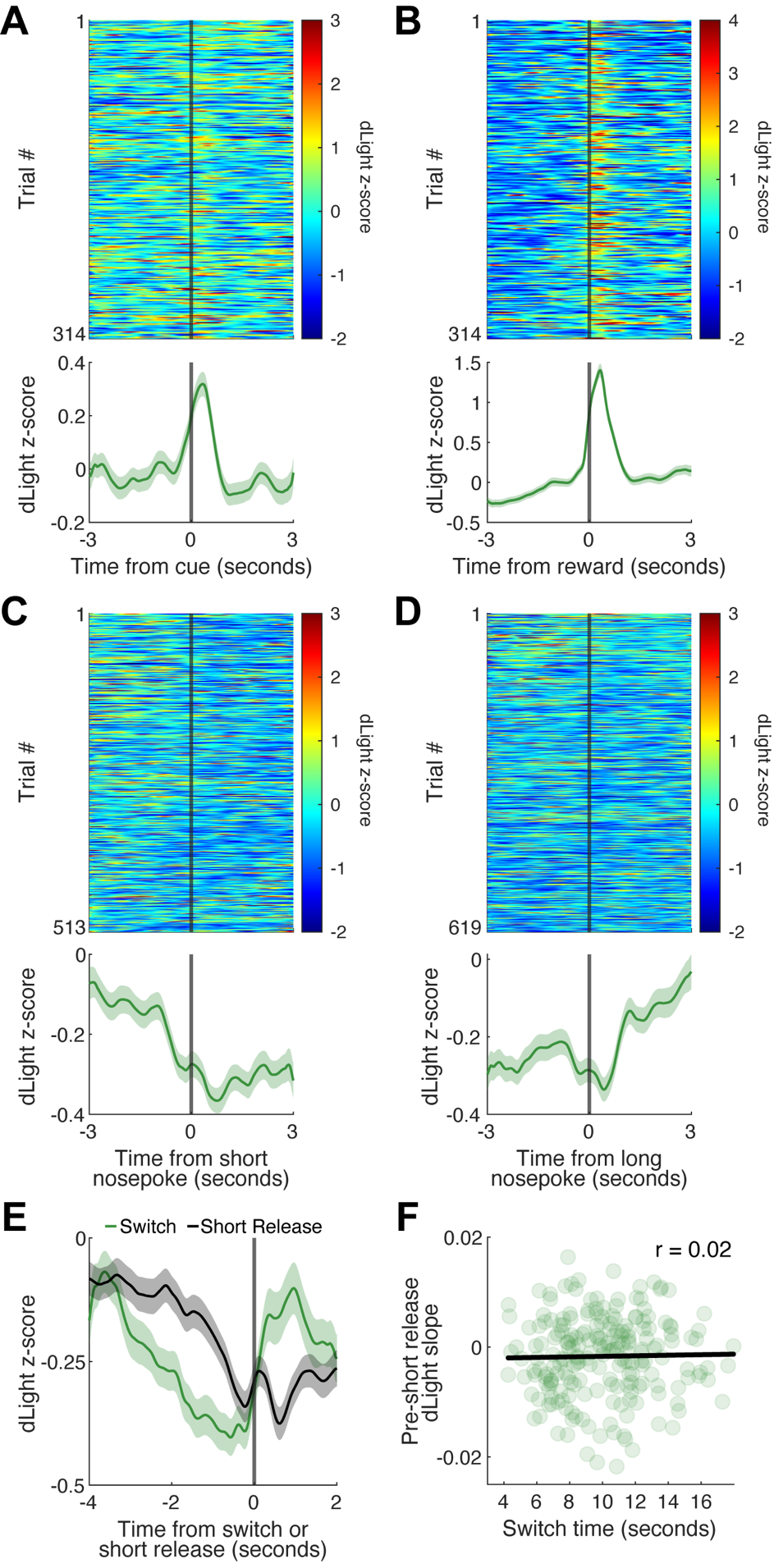

**Extended Data Figure 2.**

**A)** Top: dLight signal (z-score) aligned to cues (trial start) during every switch response trial. Trials are sorted by switch response time. Bottom: Average dLight z-score around cues. Data from 314 trials in 8 mice.

**B)** Top: dLight z-score aligned to reward delivery during every switch response trial. Trials are sorted by switch response time. Bottom: Average dLight z-score around reward delivery. Data from 314 trials in 8 mice.

**C)** Top: dLight aligned to responses at the short nosepoke during every switch response trial. Trials are sorted by time of short responses within the trial. Short responses that were the switch response and short responses within 3 seconds of trial start were removed. Bottom: Average dLight z-score around short nosepokes. Data from 314 trials in 8 mice.

**D)** Top: dLight aligned to responses at the long nosepoke during every switch response trial. Trials are sorted by time of long responses within the trial. Long responses within 3 seconds of trial end (18 seconds) were removed. Bottom: Average dLight signal (z-score) around long nosepokes. Data from 314 trials in 8 mice.

**E)** Average dLight signal aligned to the switch response (green) and average dLight signal aligned to all other short nosepoke release events (black). Data from 211 trials in 8 mice.

**F)** The slope of the dLight signal in the four second period prior to non-switch short nosepoke release events did not predict switch response time (r = 0.02, *p* = 0.78). Data from 211 trials in 8 mice.

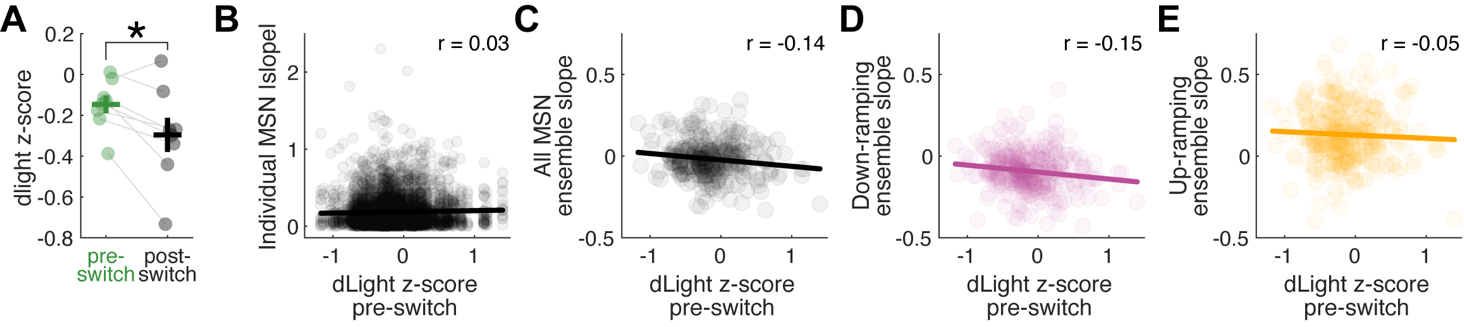

**Extended Data Figure 3.**

A) The average dLight z-score was higher in the period before the switch response than in the period after the switch response (*p* = 0.02). Horizontal lines represent group averages, vertical lines represent s.e.m., and dots represent average signal for individual mice. Data from 271 trials in 8 mice.
B) The slope of individual MSNs weakly correlated with the pre-switch dLight signal (r = 0.03, *p* = 0.04). 4106 data points from 104 MSNs in 8 mice.
C) The average ensemble slopes of all MSNs correlated with the pre-switch dLight signal (r = -0.14, *p* = 0.01). Data from 314 trials in 104 MSNs in 8 mice.
D) The average ensemble slopes of down-ramping MSNs correlated with the pre-switch dLight signal (r = -0.15 D, *p* = 0.01). Data from 314 trials in 68 MSNs in 8 mice.
E) The average ensemble slopes of up-ramping MSNs did not correlate with the pre-switch dLight signal (r = -0.05, *p* = 0.39). Data from 314 trials in 36 MSNs in 8 mice.

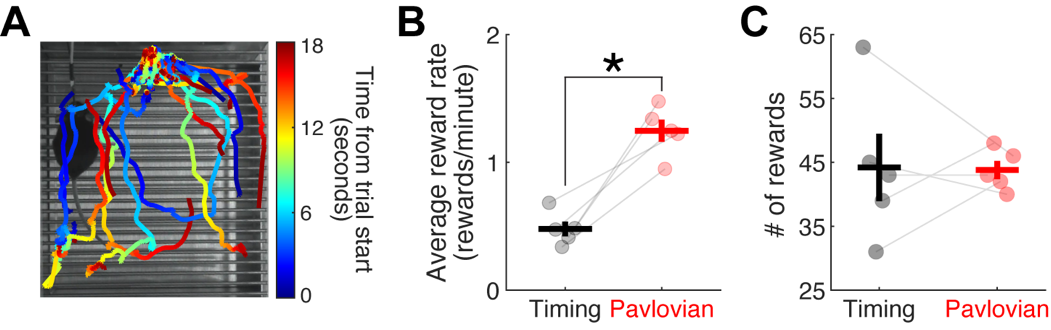

**Extended Data Figure 4.**

**A)** We tracked mouse position during Pavlovian conditioning trials. Colored lines represent the movement trajectories of a single mouse during all trials from a single session. Color represents the time within the trial aligned to trial start (seconds).

**B)** Average reward rate was higher during Pavlovian conditioning sessions (red) than interval timing sessions (black; *p* = 0.001).

**C)** Mice earned similar numbers of rewards during Pavlovian conditioning sessions (red) and interval timing sessions (black; *p* = 0.94).

Horizontal lines represent group averages, vertical lines represent s.e.m., and dots represent individual mouse averages in B-C. Data in B-C from 5 mice. **p* < 0.05

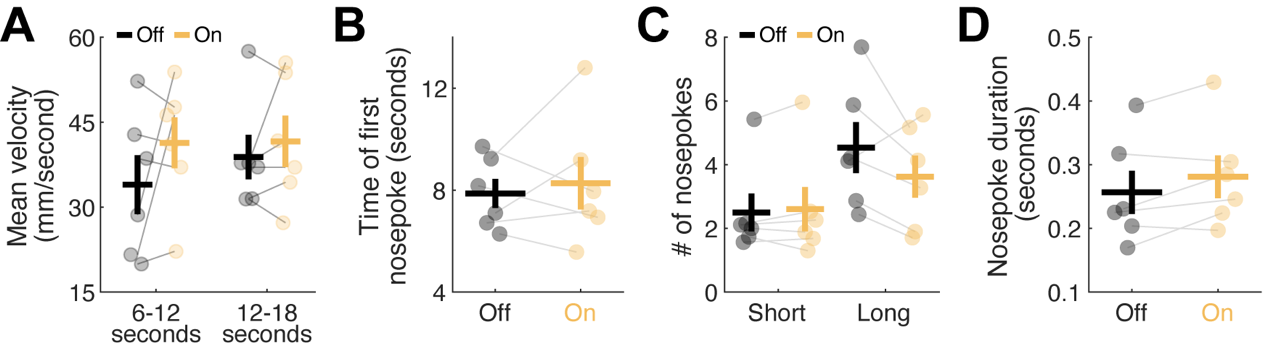

**Extended Data Figure 5.**

A) The average movement velocity of mice did not differ between laser off (black) and laser on (yellow) trials in the 6 to 12 second epoch (*p* = 0.25) or the 12 to 18 second epoch (*p* = 0.44) of the trial.
B) The time of the first nosepoke did not differ between laser off (black) and laser on (yellow) trials (*p* = 0.66).
C) The number of short and long nosepokes did not differ between laser off (black) and laser on (yellow) trials (short nosepokes: *p* = 0.49; long nosepokes: *p* = 0.16).
D) Nosepoke duration did not differ between laser off (black) and laser on (yellow) trials (*p* = 0.10).

Horizontal lines represent group averages, vertical lines represent s.e.m., and each dot represents an individual mouse average. Data from 115 laser off trials and 141 laser on trials in 6 mice.

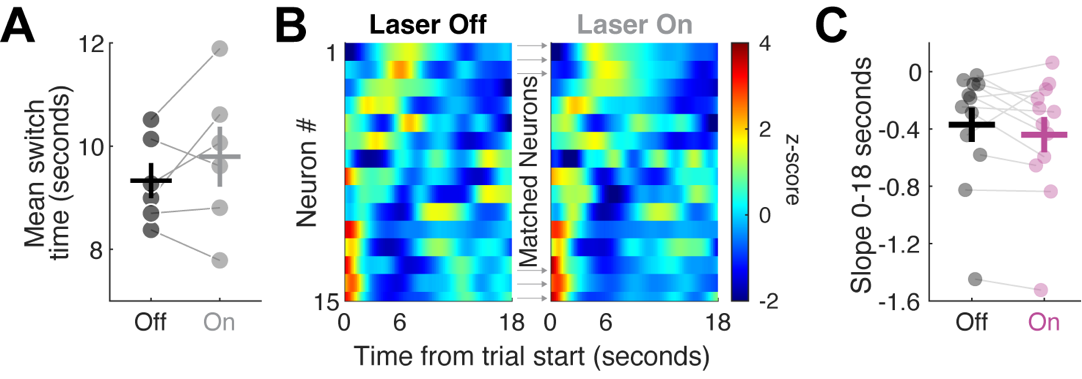

**Extended Data Figure 6.**

A) Average switch time did not differ between laser off (black) and laser on (gray) trials in control mice expressing no-opsin viruses (*p* = 0.28). Data from 330 laser off trials and 329 laser on trials in 6 mice.
B) PETHs of interval timing switch trials with the laser off (left) and with the laser on (right) in control mice. MSNs are sorted by PC1 for the laser off condition and displayed in the same order for the laser on condition. Data from 2 mice.
C) The slope of down-ramping MSNs did not change during laser on trials in control mice (*p* = 0.23). Data from 12 MSNs in 2 mice.

Horizontal lines represent group averages, vertical lines represent s.e.m., and dots represent individual mice in A and C.

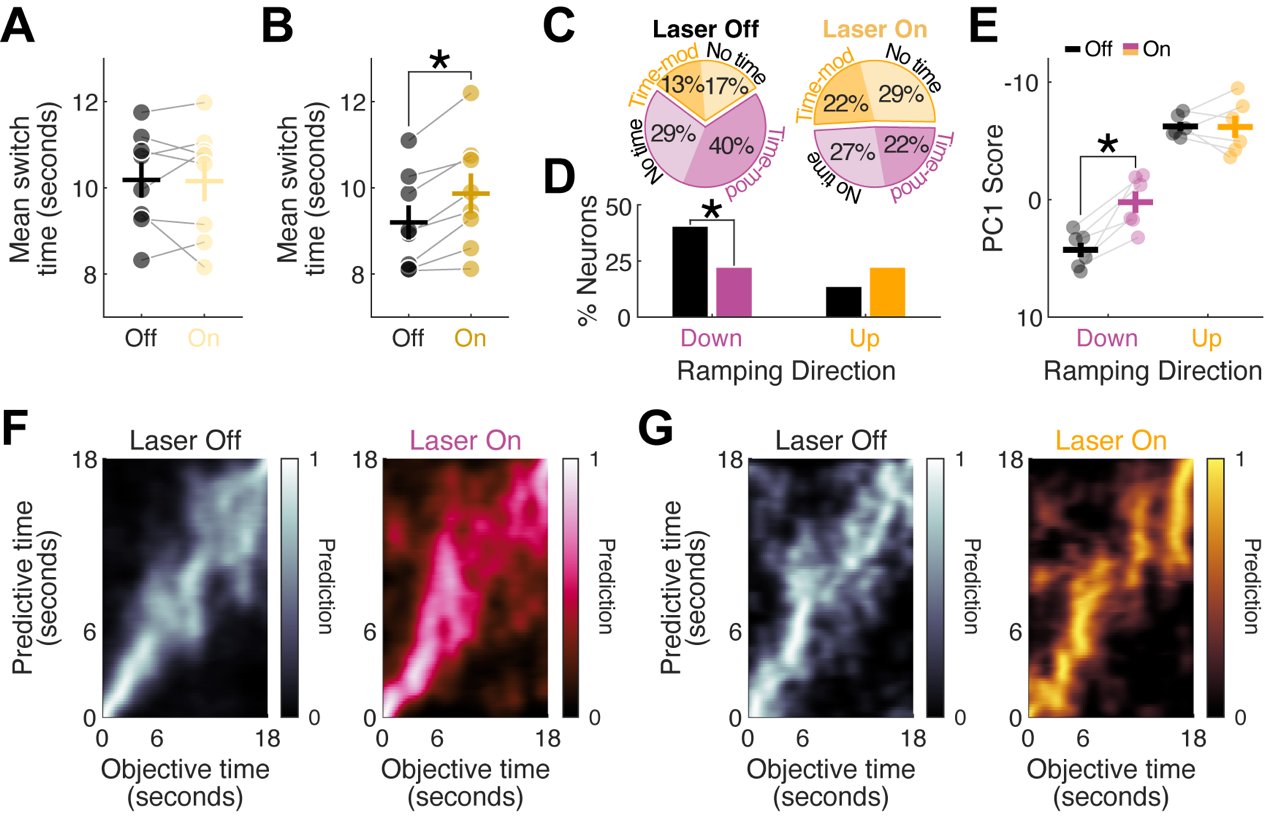

**Extended Data Figure 7.**

A) Average switch time did not differ between laser off and laser on trials in mice expressing Halorhodopsin and receiving 2 mW laser for two seconds at trial start (*p* = 0.90).
B) 5 mW laser for two seconds at trial start increased switch times in mice expressing Halorhodopsin (*p* = 0.002). Mice in A-B are the same 8 mice as in Figure 6C.
C) Percentage of MSNs that had positive slopes and GLM-defined time-modulation (dark yellow), positive slopes and no time-modulation (light yellow), negative slopes and GLM-defined time-modulation (dark pink), and negative slopes and no time-modulation (light pink). Values in pie chart are rounded to the nearest whole number.
D) Fewer down-ramping (*p* = 0.01), but not up-ramping (*p* = 0.15), MSNs had GLM-defined time-modulation during laser on trials compared to laser off trials.
E) Laser inhibition of SNc dopamine neurons decreased average ensemble PC1 scores of down-ramping MSNs (pink; *p* = 0.007) but did not affect up-ramping MSN ensembles (yellow; *p* = 0.95). Data from 6 mice (same mice as in Fig. 7). * *p* < 0.05
F) We trained a naïve Bayesian classifier to decode temporal prediction on a trial-by trial basis from firing rate from down-ramping MSN ensembles with the laser off (gray) and laser on (pink) and **G)** from up-ramping MSN ensembles with the laser off (gray) and laser on (yellow). Data from 6 mice (same data as in Fig. 7G).

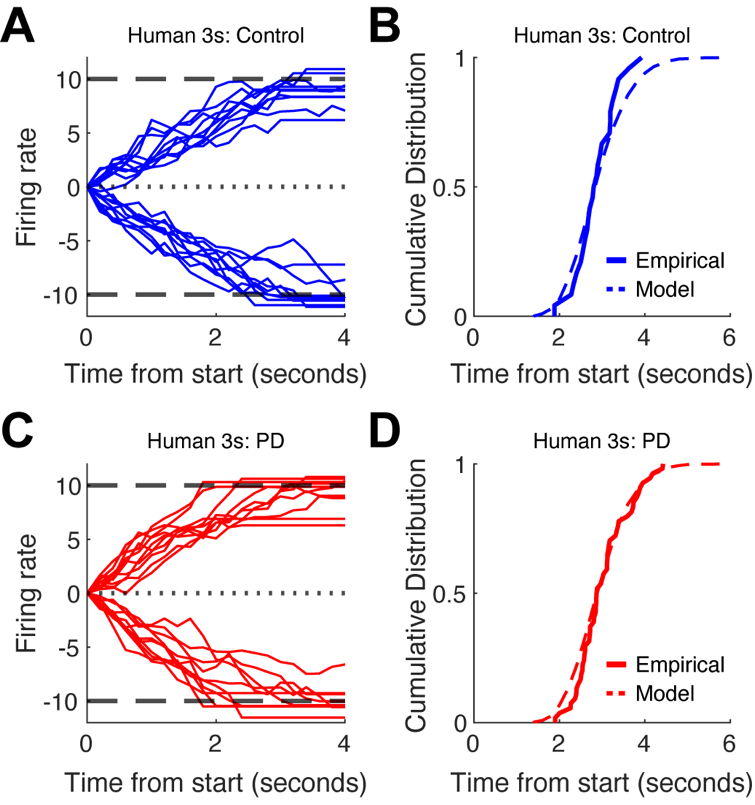

**Extended Data Figure 8.**

**A)** DDM trajectories and **B)** CDF fits for healthy control participants timing the 3 second interval.

**C)** DDM trajectories and **D)** CDF fits for PD patients timing the 3 second interval. Data from 24 control participants and 54 PD patients.
