## Supplementary Tables for "Dopamine shifts striatal ramping dynamics"

**Table 1: Experimental mice**

| **Experiments** | **Strain** | **Genotype** | **Groups** | **Number** |
| --- | --- | --- | --- | --- |
| Fiber photometry + electrophysiology | C57BL/6J | Wild-type |  | 8 (3 Female, 5 Male); only 6/8 (3 Female, 3 Male) for DeepLabCut; only 5/8 (3 Female, 2 Male) for Pavlovian conditioning |
| Fiber photometry + optogenetics | C57BL/6J | DAT-Cre |  | 2 (2 Female, 0 Male) |
| Optogenetics | C57BL/6J | DAT-Cre | Electrophysiology | 6 (3 Female, 3 Male) |
|  | C57BL/6J | DAT-Cre | Inhibition only | 2 (2 Female, 0 Male) |
|  | C57BL/6J | DAT-Cre | Control inhibition | 4 (2 Female, 2 Male) |
|  | C57BL/6J | DAT-Cre | Control inhibition + electrophysiology | 2 (1 Female, 1 Male) |

**Table 2: Table of MSNs recorded**

| **Experiments** | **Total** | **Up** | **Down** | **Up-ramping** | **Down-ramping** |
| --- | --- | --- | --- | --- | --- |
| Fiber photometry + electrophysiology |  |  |  |  |  |
| Interval timing | 104 | 36 | 68 | - | - |
| Pavlovian | 58 | - | - | - | - |
| Optogenetics |  |  |  |  |  |
| Halorhodopsin | 82 | - | - | 57 | 25 |
| Control | 15 | - | - | 12 | 3 |

**Table 3: Demographic, disease, motor, and cognitive characteristics**

|  | **Control (N=24)** | **PD (N=54)** |
| --- | --- | --- |
| Demographics and disease |  |  |
| Gender, Male/Female | 9/15 | 35/19 |
| Age (years) | 62.3 (2.6) | 69.0 (1.3) |
| Education (years) | 18.3 (0.9) | 15.7 (0.4) |
| Cognition characteristics |  |  |
| MOCA score (0 – 30) | 27.2 (0.5) | 24.6 (0.4) |
| Motor characteristics |  |  |
| mUPDRS (0 – 56) | 0.8 (0.2) | 25.1 (1.5) |

Values are expressed as mean (SEM); MOCA, Montreal Cognitive Assessment; mUPDRS, Motor Unified Parkinson’s Disease Rating Scale; PD, Parkinson’s disease.

**Table 4: Statistical Results**

| Figure | Comparison | Statistical Test | n | Test Statistic | r | Confidence Interval (s.e.m.) | p-value |
| --- | --- | --- | --- | --- | --- | --- | --- |
| Figure 2B | 0-18 seconds All MSNs vs. Down MSNs | *lm* w/ post-hoc comparisons | 20 | 0.43 |  |  | 0.99 |
| Figure 2B | 0-18 seconds All MSNs vs. Up MSNs | *lm* w/ post-hoc comparisons | 20 | 1.93 |  |  | 0.39 |
| Figure 2B | 0-18 seconds Down MSNs vs. Up MSNs | *lm* w/ post-hoc comparisons | 20 | 1.49 |  |  | 0.67 |
| Figure 2B | 0-18 seconds All MSNs vs. Shuffled All | *lm* w/ post-hoc comparisons | 20 | 12.74 |  |  | **<0.0001** |
| Figure 2B | 0-18 seconds Down MSNs vs. Shuffled Down | *lm* w/ post-hoc comparisons | 20 | 12.39 |  |  | **<0.0001** |
| Figure 2B | 0-18 seconds Up MSNs vs. Shuffled Up MSNs | *lm* w/ post-hoc comparisons | 20 | 10.69 |  |  | **<0.0001** |
| Figure 2B | 0-6 seconds All MSNs vs. Down MSNs | *lm* w/ post-hoc comparisons | 20 | 0.11 |  |  | 1.0 |
| Figure 2B | 0-6 seconds All MSNs vs. Up MSNs | *lm* w/ post-hoc comparisons | 20 | 1.48 |  |  | 0.67 |
| Figure 2B | 0-6 seconds Down MSNs vs. Up MSNs | *lm* w/ post-hoc comparisons | 20 | 1.38 |  |  | 0.74 |
| Figure 2B | 0-6 seconds All MSNs vs. Shuffled All | *lm* w/ post-hoc comparisons | 20 | 9.19 |  |  | **<0.0001** |
| Figure 2B | 0-6 seconds Down MSNs vs. Shuffled Down | *lm* w/ post-hoc comparisons | 20 | 9.44 |  |  | **<0.0001** |
| Figure 2B | 0-6 seconds Up MSNs vs. Shuffled Up | *lm* w/ post-hoc comparisons | 20 | 6.81 |  |  | **<0.0001** |
| Figure 2C | Individual MSN slope vs. Switch time | Pearson linear correlation | 3904 |  | 0.05 | [0.02, 0.08] | **0.003** |
| Figure 2D | Ensemble MSN slope vs. Switch time | Pearson linear correlation | 300 |  | 0.12 | [0.001, 0.23] | **0.047** |
| Figure 2E | Ensemble down MSN slope vs. Switch time | Pearson linear correlation | 300 |  | 0.14 | [0.03, 0.25] | **0.01** |
| Figure 2F | Ensemble up MSN slope vs. Switch time | Pearson linear correlation | 300 |  | 0.06 | [-0.05, 0.17] | 0.30 |
| Figure 3B | 2 seconds: laser on vs. off | Wilcoxon rank sum | 21 | 3.87 |  |  | **<0.0001** |
| Figure 3B | 7 seconds: laser on vs. off | Wilcoxon rank sum | 21 | 4.70 |  |  | **<0.0001** |
| Figure 3H | Switch time vs. Pre-switch dLight | Pearson linear correlation | 273 |  | 0.02 | [-0.10, 0.14] | 0.77 |
| Figure 3I | Switch time vs. Pre-switch dLight slope | Pearson linear correlation | 276 |  | 0.24 | [0.12, 0.35] | **<0.0001** |
| Figure 3K | Correlation 0-18s vs. Shuffled | Wilcoxon rank sum | 6 | 57.00 |  |  | **0.002** |
| Figure 3K | Correlation 0-6s vs. Shuffled | Wilcoxon rank sum | 6 | 57.00 |  |  | **0.002** |
| Figure 3L | Pre-switch dLight slope vs. velocity slope pre-switch | Pearson linear correlation | 191 |  | 0.08 | [-0.06, 0.22] | 0.26 |
| Extended Data Figure 2F | Switch time vs. non-switch slope | Pearson linear correlation | 191 |  | 0.02 | [-0.12, 0.15] | 0.78 |
| Figure 4C | Bayes 0-18s MSNs vs. dLight | *lm* w/ post-hoc comparisons | 20 | 18.49 |  |  | **<0.0001** |
| Figure 4C | Bayes 0-18s MSNs vs. shuffled MSNs | *lm* w/ post-hoc comparisons | 20 | 16.55 |  |  | **<0.0001** |
| Figure 4C | Bayes 0-18s dLight vs. shuffled dLight | *lm* w/ post-hoc comparisons | 20 | 3.28 |  |  | **8.5×10^-3^** |
| Figure 4C | Bayes 0-6s MSNs vs. dLight | *lm* w/ post-hoc comparisons | 20 | 30.38 |  |  | **<0.0001** |
| Figure 4C | Bayes 0-6s MSNs vs. shuffled MSNs | *lm* w/ post-hoc comparisons | 20 | 17.15 |  |  | **<0.0001** |
| Figure 4C | Bayes 0-6s dLight vs. shuffled dLight | *lm* w/ post-hoc comparisons | 20 | 1.34 |  |  | 0.54 |
| Figure 4F | -6 to 0 seconds – Down MSNs | LMM; contrasts vs. 0 | 8 | 3.33 |  |  | **0.01** |
| Figure 4F | 0 to 6 seconds – Down MSNs | LMM; contrasts vs. 0 | 8 | 5.03 |  |  | **<0.0001** |
| Figure 4F | 6 to 12 seconds – Down MSNs | LMM; contrasts vs. 0 | 8 | 4.64 |  |  | **1.3×10^-4^** |
| Figure 4F | 12 to 18 seconds – Down MSNs | LMM; contrasts vs. 0 | 8 | 4.72 |  |  | **1.0×10^-4^** |
| Figure 4F | 18 to 24 seconds – Down MSNs | LMM; contrasts vs. 0 | 8 | 2.38 |  |  | 0.08 |
| Figure 4F | -6 to 0 seconds – Up MSNs | LMM; contrasts vs. 0 | 8 | -0.74 |  |  | 0.92 |
| Figure 4F | 0 to 6 seconds – Up MSNs | LMM; contrasts vs. 0 | 8 | 1.53 |  |  | 0.40 |
| Figure 4F | 6 to 12 seconds – Up MSNs | LMM; contrasts vs. 0 | 8 | 3.33 |  |  | **0.01** |
| Figure 4F | 12 to 18 seconds – Up MSNs | LMM; contrasts vs. 0 | 8 | 3.05 |  |  | **0.02** |
| Figure 4F | 18 to 24 seconds – Up MSNs | LMM; contrasts vs. 0 | 8 | 0.65 |  |  | 0.92 |
| Figure 4G | dLight z-score 0 to 6s vs. 6 to 12s | Paired t-test | 8 | 10.50 |  |  | **<0.0001** |
| Figure 4H | Individual MSN slope vs. dLight z-score 0-6s | Pearson linear correlation | 4106 |  | 0.03 | [0.003, 0.06] | **0.03** |
| Figure 4I | Ensemble MSN slope vs. dLight z-score 0-6s | Pearson linear correlation | 314 |  | -0.17 | [-0.28,-0.06] | **0.002** |
| Figure 4J | Down MSN ensemble slope vs. dLight z-score 0-6s | Pearson linear correlation | 314 |  | -0.20 | [-0.30,-0.09] | **4.7×10^-4^** |
| Figure 4K | Up MSN ensemble slope vs. dLight z-score 0-6s | Pearson linear correlation | 314 |  | -0.05 | [-0.16, 0.06] | 0.33 |
| Extended Data Figure 3A | dLight pre- vs. post-switch | Paired t-test | 8 | 3.12 |  |  | **0.02** |
| Extended Data Figure 3B | Individual MSN slope vs. dLight z-score pre-switch | Pearson linear correlation | 4106 |  | 0.03 | [0.002, 0.06] | **0.04** |
| Extended Data Figure 3C | Ensemble MSN slope vs. dLight z-score pre-switch | Pearson linear correlation | 314 |  | -0.14 | [-0.25, 0.03] | **0.01** |
| Extended Data Figure 3D | Down MSN slope vs. dLight z-score pre-switch | Pearson linear correlation | 314 |  | -0.15 | [-0.26, 0.04] | **0.01** |
| Extended Data Figure 3E | Up MSN slope vs. dLight z-score pre-switch | Pearson linear correlation | 314 |  | -0.05 | [-0.16, 0.06] | 0.39 |
| Figure 5C | % Ramping Timing vs. Pavlov | Chi-squared test |  | 10.44 |  |  | **0.001** |
| Figure 5C | % Ramping Timing vs. Both | Chi-squared test |  | 12.94 |  |  | **0.0003** |
| Figure 5C | % Ramping Pavlov vs. Both | Chi-squared test |  | 0.34 |  |  | 0.56 |
| Figure 5D | MSNs trial start: timing vs. Pavlov | Paired t-test | 5 | -1.47 |  |  | 0.22 |
| Figure 5D | MSNs reward: timing vs. Pavlov | Paired t-test | 5 | -1.56 |  |  | 0.19 |
| Figure 5E | MSNs slope: timing vs. Pavlov | Paired t-test | 5 | 5.59 |  |  | **0.005** |
| Figure 5H | dLight trial start: timing vs. Pavlov | Paired t-test | 5 | 0.85 |  |  | 0.44 |
| Figure 5H | dLight reward: timing vs. Pavlov | Paired t-test | 5 | 0.27 |  |  | 0.80 |
| Figure 5I | dLight slope: timing vs. Pavlov | Paired t-test | 5 | -2.38 |  |  | 0.08 |
| Figure 5J | dLight z-score trial start vs. ensemble slope | Pearson linear correlation | 219 |  | 0.04 | [-0.09, 0.18] | 0.51 |
| Figure 5K | dLight z-score trial start vs. ensemble trial start | Pearson linear correlation | 219 |  | 0.01 | [-0.12, 0.14] | 0.91 |
| Figure 5L | dLight z-score reward vs. ensemble reward | Pearson linear correlation | 219 |  | 0.15 | [0.01, 0.27] | **0.03** |
| Extended Data Figure 4B | Average reward rate: timing vs. Pavlov | Paired t-test | 5 | -8.28 |  |  | **0.001** |
| Extended Data Figure 4C | # rewards: timing vs. Pavlov | Paired t-test | 5 | 0.08 |  |  | 0.94 |
| Figure 6C | Switch time: Laser off vs. on | Paired t-test | 9 | -2.83 |  |  | **0.02** |
| Figure 6D | Velocity 0-18s Laser off vs. on | Paired t-test | 6 | -1.37 |  |  | 0.23 |
| Figure 6D | Velocity 0-6s Laser off vs. on | Paired t-test | 6 | -1.24 |  |  | 0.27 |
| Extended Data Figure 5A | Velocity 6-12s Laser off vs. on | Paired t-test | 6 | -1.31 |  |  | 0.25 |
| Extended Data Figure 5A | Velocity 12-18s Laser off vs. on | Paired t-test | 6 | -0.84 |  |  | 0.44 |
| Extended Data Figure 5B | First nosepoke: off vs. on | Paired t-test | 6 | -0.47 |  |  | 0.66 |
| Extended Data Figure 5C | # Short NP: off vs. on | Paired t-test | 6 | -0.74 |  |  | 0.49 |
| Extended Data Figure 5C | # Long NP: off vs. on | Paired t-test | 6 | 1.67 |  |  | 0.16 |
| Extended Data Figure 5D | NP Duration: off vs. on | Paired t-test | 6 | -2.04 |  |  | 0.10 |
| Figure 7C | % Ramping Down Laser on vs. off | Chi-squared test |  | 7.29 |  |  | **0.01** |
| Figure 7D | Slope down: laser off vs. on | Paired t-test | 57 | -4.54 |  |  | **<0.0001** |
| Figure 7D | Slope up: laser off vs. on | Paired t-test | 25 | -1.15 |  |  | 0.26 |
| Figure 7F | PC1 down: laser off vs. on | Paired t-test | 57 | 6.67 |  |  | **<0.0001** |
| Figure 7F | PC1 up: laser off vs. on | Paired t-test | 25 | 1.94×10^-4^ |  |  | 1.00 |
| Figure 7G | Bayes Down: Off vs. On | Wilcoxon rank sum | 20 | 3.50 |  |  | **0.0005** |
| Figure 7G | Bayes Up: Off vs. On | Wilcoxon rank sum | 20 | -0.55 |  |  | 0.58 |
| Extended Data Figure 6A | Switch time: Laser off vs. on | Paired t-test | 6 | -1.20 |  |  | 0.28 |
| Extended Data Figure 6C | Slope down: laser off vs. on | Paired t-test | 6 | 1.26 |  |  | 0.23 |
| Extended Data Figure 7A | Switch time: Laser off vs. on | Paired t-test | 8 | 0.12 |  |  | 0.90 |
| Extended Data Figure 7B | Switch time: Laser off vs. on | Paired t-test | 8 | -4.94 |  |  | **0.002** |
| Extended Data Figure 7D | % Time-modulated Down | Chi-squared test |  | 6.40 |  |  | **0.01** |
| Extended Data Figure 7D | % Time-modulated Up | Chi-squared test |  | 2.05 |  |  | 0.15 |
| Extended Data Figure 7E | PC1 down-ramping ensemble: Laser off vs. on | Paired t-test | 6 | 4.39 |  |  | **0.007** |
| Extended Data Figure 7E | PC1 up-ramping ensemble: Laser off vs. on | Paired t-test | 6 | -0.06 |  |  | 0.95 |
| Figure 8G | Main effect of disease | Linear mixed-effects model | 153 | 7.43 |  |  | **0.007** |
| Figure 8G | Main effect of interval | Linear mixed-effects model | 153 | 284.49 |  |  | **<0.001** |
| Figure 8G | Disease-by-interval interaction | Linear mixed-effects model | 153 | 0.41 |  |  | 0.51 |
| Figure 8G | Main effect of age | Linear mixed-effects model | 153 | 7.20 |  |  | **0.008** |
| Figure 8G | Main effect of disease (age model) | Linear mixed-effects model | 153 | 3.57 |  |  | **0.06** |
| Figure 8G | Main effect of sex | Linear mixed-effects model | 153 | 3.17 |  |  | 0.08 |
| Figure 8G | Main effect of disease (sex model) | Linear mixed-effects model | 153 | 4.89 |  |  | **0.03** |
| Figure 8G | Main effect of education | Linear mixed-effects model | 153 | 2.21 |  |  | 0.14 |
| Figure 8G | Main effect of disease (disease model) | Linear mixed-effects model | 153 | 4.18 |  |  | **0.04** |
